# CryoEM Structures of the Nitrogenase Complex During Catalytic Turnover

**DOI:** 10.1101/2022.06.05.494884

**Authors:** Hannah L. Rutledge, Brian D. Cook, Hoang P. M. Nguyen, Mark A. Herzik, Akif Tezcan

**Affiliations:** Department of Chemistry and Biochemistry, University of California, San Diego, La Jolla, CA 92093, USA

## Abstract

The enzyme nitrogenase couples adenosine triphosphate (ATP) hydrolysis to the multi-electron reduction of atmospheric dinitrogen into ammonia. Despite extensive research, the mechanistic details of ATP-dependent energy transduction and dinitrogen reduction by nitrogenase are not well understood, requiring new strategies to monitor its structural dynamics during catalytic action. Here we report the cryogenic electron microscopic interrogation of the nitrogenase complex under enzymatic turnover conditions, which has enabled the structural characterization of the nitrogenase reaction intermediates at high resolution for the first time. Our structures show that asymmetry governs all aspects of nitrogenase mechanism including ATP hydrolysis, protein-protein interactions, and catalysis. Furthermore, they reveal several previously unobserved, mechanistically relevant conformational changes near the catalytic iron-molybdenum cofactor that are correlated with the nucleotide-hydrolysis state of the enzyme.

**One-sentence summary:** High-resolution cryoEM structures of the nitrogenase complex obtained under turnover provide new mechanistic insights.

## Main text

Reduced forms of nitrogen are essential for the biosynthesis of aminoand nucleic acids as well as the production of fertilizers and many commodity chemicals^(1)^. As the only enzyme capable of nitrogen fixation, nitrogenase catalyzes the eight-electron reduction of atmospheric nitrogen (N_2_) and protons (H^+^) into ammonia (NH_3_) and hydrogen (H_2_)^(2, 3)^ **(Fig. 1A)**. Nitrogenase is a two-component enzyme, which, in its most common form, consists of the iron protein (FeP, *γ*_2_ homodimer) and the molybdenum-iron protein (MoFeP, *α*_2_ *β* _2_ heterotetramer) **(Fig. 1B)**^(4, 5)^. Alongside its unique capability of N_2_ fixation, nitrogenase is distinct from most redox enzymes in its requirement for adenosine triphosphate (ATP) hydrolysis to enable the successive transfer of electrons and protons for substrate reduction^(6, 7)^. The coupling of ATP hydrolysis to electron transfer (ET) is mediated by FeP, which forms a specific, nucleotide-dependent complex with MoFeP, and hydrolyzes two ATP molecules for the transfer of an electron to MoFeP^(4-7)^. This part of nitrogenase catalysis is termed the “FeP cycle” **(Fig. 1A)**^(2, 8)^. In the “MoFeP cycle” **(Fig. 1A)**^(2, 8)^, the electrons from FeP are received by the P-cluster (an [8Fe:7S] complex) of MoFeP and relayed to the iron-molybdenum cofactor (FeMoco, a [7Fe:9S:C:Mo]-homocitrate complex) where N_2_ binding and reduction occur. Specific ATP-dependent interactions between FeP and MoFeP are necessary not only for interprotein ET, but also for gated ET between the P-cluster and FeMoco through long-distance conformational perturbations, the nature of which are not understood^(6, 7)^.

**Figure 1.**
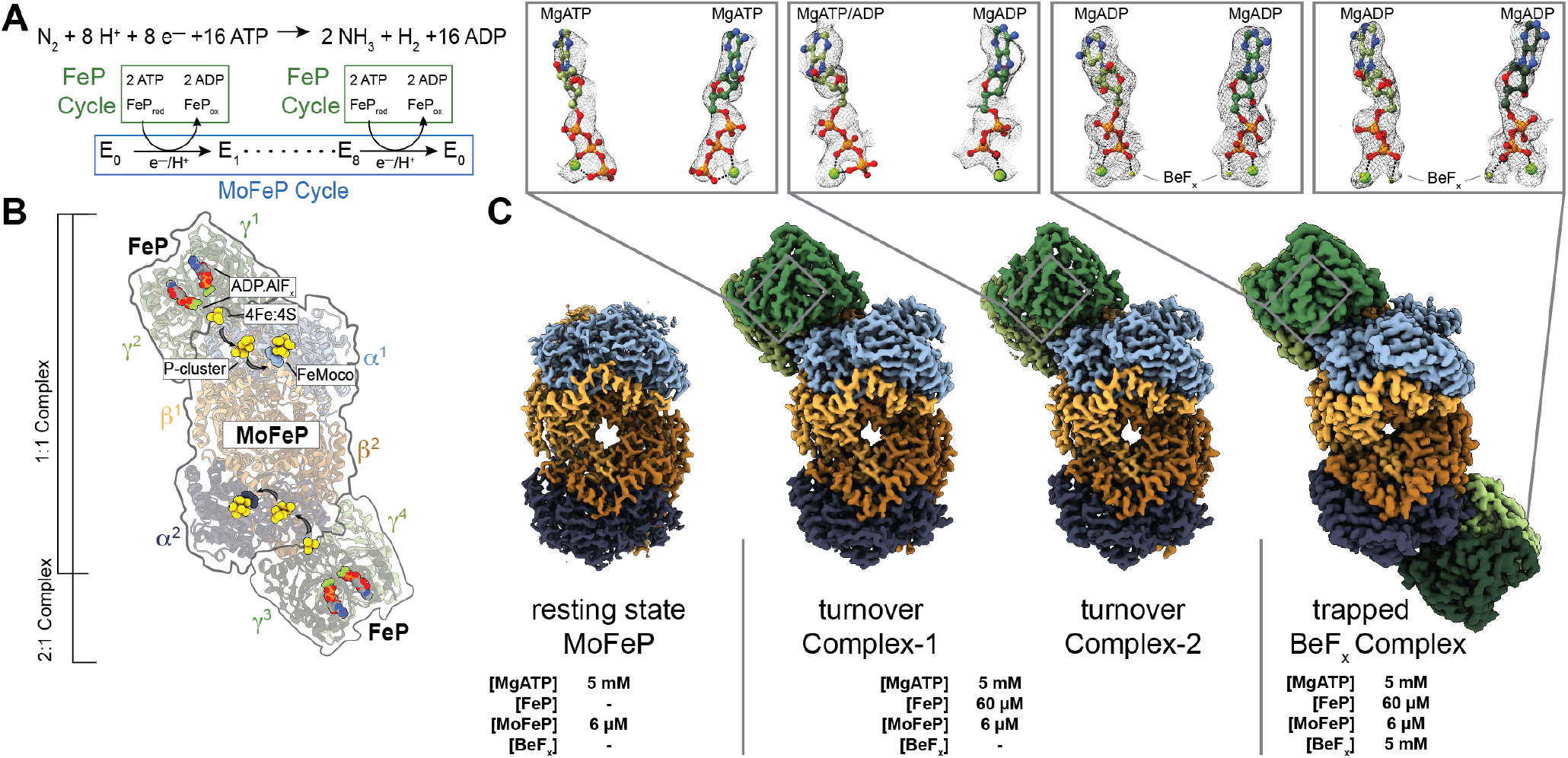
Catalytic cycle for nitrogenase and its structural characterization under non-turnover and turnover conditions. **(A)** Chemical reaction catalyzed by nitrogenase. The FeP and MoFeP cycles involved in this reaction are shown in green and blue boxes, respectively. There are eight FeP cycles in each MoFeP cycle. **(B)** Crystal structure (PDB ID: 1M34) of the 2:1 FeP:MoFeP complex stabilized by MgADP.AlF_x_, showing the relative positions of the individual FeP (*γ*^1^ and *γ*^2^ and *γ*^3^ and *γ*^4^) and MoFeP (*α*^1^*β*^1^*α*^2^*β*^2^) subunits, the nucleotides, and the metalloclusters. The FeP subunits are shown in dark green (*γ*^1^ or *γ*^3^) and light green (*γ*^2^ or *γ*^4^), and the MoFeP subunits are highlighted in light blue (*α*^1^), dark blue (*α*^2^), light orange (*β* ^1^) and dark orange (*β* ^2^). Black arrows indicate the path of electron flow. **(C)** CryoEM maps of resting state MoFeP (^rs^MoFeP; ∼1.8 Å resolution) obtained in the absence of FeP, and the turnover Complex-1 (^t/o^Complex-1; ∼2.3 Å resolution) and turnover Complex-2 (^t/o^Complex-2; ∼2.3 Å resolution) obtained under turnover conditions, and BeF_x_ trapped FeP-MoFeP complex (∼2.4 Å resolution). Coloring scheme for the subunits is the same as in **(B)**. Magnified views of the nucleotides bound to the nitrogenase complexes and their corresponding cryoEM densities are shown in boxes.

While the dynamic coupling between ATP hydrolysis and ET ultimately drives N_2_ fixation, it also creates experimental challenges for a mechanistic understanding of nitrogenase. First, substrates and inhibitors can only bind reduced forms of FeMoco, whose generation requires continuous turnover conditions that include ATP and reduced FeP^(2, 3)^. Because MoFeP is also an inherent hydrogenase, such reduced forms of FeMoco promptly return to the resting state through H_2_ evolution upon termination of ATP hydrolysis^(8, 9)^, rendering substrate-bound states of FeMoco too fleeting for structural characterization. Second, the necessity of continuous ATP hydrolysis for catalysis leads to a heterogeneous distribution of nitrogenase sub-states which are difficult to interrogate experimentally. These sub-states differ not only in the FeP-MoFeP complexation state and the extent of ATP hydrolysis within the FeP cycle, but also in the oxidation states of the metal cofactors and the reaction intermediates on FeMoco (E_0_-E_8)_ within the MoFeP cycle **(Fig. 1A-B)**.

Previous crystallographic studies have provided detailed views of the nucleotide-dependent conformations of the FeP-MoFeP complexes and outlined the path of ET in nitrogenase **(Fig. 1B)**^(10, 11)^. Yet, these static views – obtained under non-turnover conditions and constrained by crystal lattice packing – showed no variation in the structure of MoFeP, providing little insight into the mechanism of ATP/FeP-mediated redox events within this protein. Recent crystallographic studies of MoFeP and its vanadium analog, VFe-protein, revealed possible modes of ligand interactions with FeMoco and showed that the cofactor is capable of undergoing compositional changes^(12-14)^. However, these structural snapshots were obtained using crystals that formed hours to days after the last enzymatic turnover reaction and their catalytic relevance is an open question. In parallel, extensive freeze-quench spectroscopic studies of nitrogenase have characterized substrate- and intermediate-bound states of FeMoco^(15, 16)^. Yet, these methods do not report on the atomic structure of the cofactor as a whole and provide little information regarding how the local cofactor-ligand interactions are linked to global ATP/FeP-dependent structural dynamics of MoFeP. Consequently, we lack a detailed understanding of why and how ATP hydrolysis is used to drive N_2_ fixation and how catalysis at FeMoco proceeds. Clearly, new experimental approaches are needed to structurally interrogate nitrogenase during catalysis at or near atomic resolution to simultaneously characterize the ATP-dependent FeP-MoFeP interactions and the associated structural dynamics within each protein.

We sought to overcome the limitations of prior studies by using cryogenic electron microscopy (cryoEM) to directly visualize nitrogenase under enzymatic turnover conditions. We prepared cryoEM samples of *Azotobacter vinelandii* nitrogenase under high-electron flux turnover conditions to maximize NH_3_ production. The optimized samples contained a 10-fold molar excess of FeP over MoFeP and low ionic strength (≤25 mM NaCl) to favor the formation of electrostatically driven FeP-MoFeP complexes. We included moderately high MgATP and reductant (dithionite) concentrations (both 5 mM) to ensure that they are not depleted during turnover while minimizing background electron scattering. Samples were prepared anaerobically under a N_2_ atmosphere and immediately flash frozen in liquid N_2_ following initiation of turnover. The 30-s sample preparation period was sufficiently long to ensure that steady-state catalytic conditions were reached but short enough such that there was still excess reductant in solution and no significant MgADP build-up. We collected a large cryoEM dataset (*>*15K movies) which yielded *>*4.5 million usable particles **(figs. S1, S2; table S1)**. Through exhaustive 2-and 3-D classification and refinement, we isolated free MoFeP (∼60%) and FeP particles (∼5%) as well as FeP-MoFeP nitrogenase assemblies (∼35%) from this heterogeneous mixture. We determined the structures of two conformationally distinct 1:1 FeP:MoFeP complexes under turnover, designated “^t/o^Complex-1” and “^t/o^Complex-2”, at ∼2.3 Å resolution **(Fig. 1C)**. As a reference, we also obtained ∼1.8-Å resolution cryoEM structure of resting-state MoFeP, termed “rsMoFeP”, using the same turnover conditions but in the absence of FeP **(Fig 1C; fig. S3, table S2)**.

Given the *C*_*2*_ symmetry of MoFeP and the large separation (*>*65 Å) between the nearest clusters from the symmetry-related *αβ* subunits, it has long been assumed that the two *αβ* halves of MoFeP function independently from one another^(2, 8)^. In support of this assumption, the crystal structures of various FeP-MoFeP complexes in different nucleotide-bound states largely possess a 2:1 FeP:MoFeP stoichiometry **(Fig. 1B)**^(10, 11, 17)^. Unexpectedly, our turnover samples did not contain any particles that could be assigned to a 2:1 FeP:MoFeP complex **(Fig. 1C; fig. S1)**. We considered the possibility that the exclusive observation of 1:1 complexes in our turnover samples could arise from an experimental artifact such as protein degradation or increase in ionic strength during cryoEM grid preparation. Therefore, we prepared a second set of turnover samples as above that also included 5 mM beryllium fluoride (BeF_x_), which is known to arrest ATP hydrolysis in a transition-like state to yield quasi-irreversible, solution-stable 2:1 and 1:1 FeP-MoFeP complexes^(18)^. Accordingly, our cryoEM samples contained a large fraction of 2:1 FeP:MoFeP complexes alongside 1:1 species, but no detectable free MoFeP particles **(figs. S4, S5; table S3)**. We determined the structure of the MgADP.BeF_x_-bound 2:1 FeP:MoFeP at ∼2.4 Å resolution and found it to be isostructural to the crystal structure of the related MgADP.AlF_x_-bound 2:1 FeP:MoFeP complex (PDB ID: 1M34; **Fig. 1C, fig. S5**). These observations affirm that our cryoEM samples contain intact proteins and operate under native turnover conditions, in turn indicating that the 1:1 FeP:MoFeP stoichiometry is the predominant nitrogenase assembly state during catalysis.

Prior work using pre-steady-state kinetics measurements revealed that the extents of interprotein ET and ATP hydrolysis were approximately half of what would be expected if there were two independent FeP binding sites on MoFeP^(19)^. Originally, such half-reactivity was attributed to either partial inactivity of FeP molecules^(19)^ or to the possible existence of an alternative interaction mode between FeP and MoFeP^(20)^. Recent studies favored a model of negative cooperativity within a 2:1 FeP:MoFeP complex, whereby one of the bound FeP molecules suppresses ATP hydrolysis by the other bound FeP and the redox activity of the opposite *αβ* half of MoFeP^(21, 22)^. Our cryoEM structures instead suggest that half-reactivity and negative cooperativity in nitrogenase arise from MoFeP binding to only one FeP molecule at a time during turnover.

^t/o^Complex-1 and ^t/o^Complex-2 were distinguished during cryoEM data processing based primarily on the structural variability of the FeP components. Thus, we first examined whether these differences are associated with the ATP-hydrolysis state of the two complexes. Earlier crystal structures identified at least three nucleotide-state-dependent FeP-MoFeP docking geometries (DG1, DG2 and DG3) and led to the hypothesis that FeP moves in a unidirectional fashion across the MoFeP surface during turnover **(Fig. 2A)**^(6, 11)^. The DG1 state predominates in the absence of nucleotides but is also populated in the presence of ATP^(23)^ and corresponds to an electrostatically guided encounter complex wherein FeP largely interacts with the *β* -subunit of MoFeP. DG2 is the activated nitrogenase complex in which ATP hydrolysis is coupled to interprotein ET, with FeP occupying the quasi-symmetric surface of MoFeP shared between *α* and *β* subunits **(Figs. 1B, 2A)**^(10, 11)^. Finally, DG3 is formed by ADP-bound FeP and primarily utilizes the *α*-subunit surface of MoFeP^(11)^. The cryoEM analysis of our turnover samples revealed that ^t/o^Complex-1 and ^t/o^Complex-2 were exclusively in DG2 **(Fig.2A)**, implying that this configuration has a higher stability and/or longer residence time relative to DG1 and DG3. Characteristic of a DG2 configuration, both complexes feature extensive interactions between FeP and MoFeP (buried surface areas *>*3600 Å^2^) and a short [4Fe:4S]-to-P-cluster edge-toedge distance of 15 Å, primed for rapid interprotein ET **(Fig. 2B)**. In ^t/o^Complex-1, both FeP *γ* subunits (*γ*^1^ and *γ*^2^) are occupied by ATP molecules with clear densities for the *γ*-phosphate groups and associated Mg^2+^ ions **(Fig. 1C)**. By contrast, the *γ*^1^ subunit of ^t/o^Complex-2 features an ATP molecule with weak density for the *γ*-phosphate, whereas the *γ*^2^ subunit is ADP-bound and the *γ*-phosphate is completely absent from the nucleotide binding pocket, indicative of asymmetry in ATP hydrolysis **(Fig. 1C)**. This observation is consistent with the crystal structure of a mixed-nucleotide FeP-MoFeP complex, in which AMPPCP (a non-hydrolyzable ATP analog) and ADP were selectively bound to the *γ*^1^ and *γ*^2^ subunits, respectively^(24)^. The differences in the nucleotide occupancies of ^t/o^Complex-1 and ^t/o^Complex-2 are reflected in their distinct FeP conformations **(Fig. 2C)**, which is further corroborated by a principal component analysis of available FeP structures **(fig. S6)**. Collectively, these observations indicate that (1) our cryoEM samples represent active turnover conditions, (2) the hydrolysis of two ATP molecules in each FeP cycle occurs in a stepwise fashion, and (3) ^t/o^Complex-1 and ^t/o^Complex-2 correspond, respectively, to preand midATP hydrolysis states of the nitrogenase complex that are populated along the catalytic reaction coordinate.

**Figure 2.**
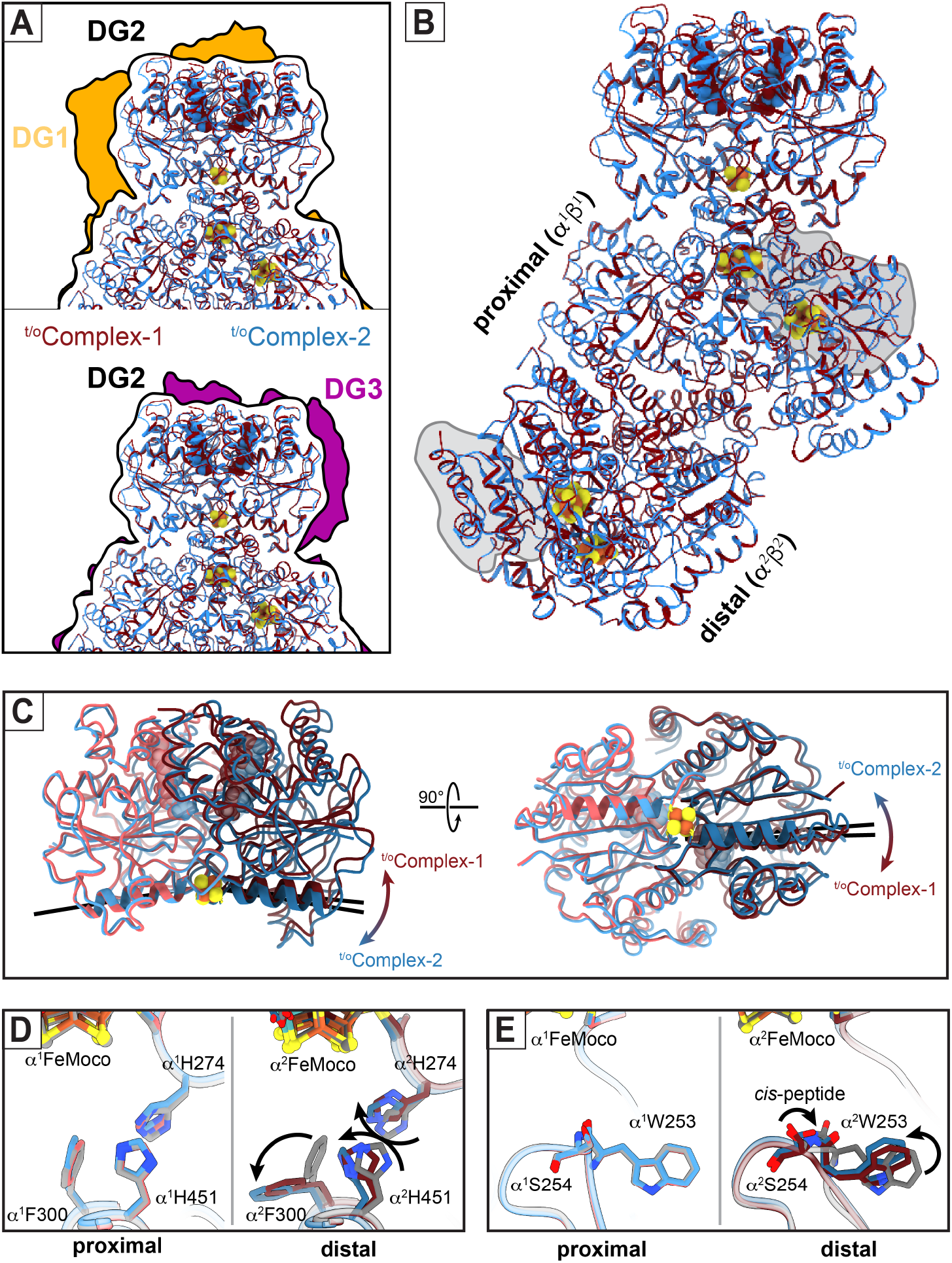
Structural details of nitrogenase complexes characterized under turnover conditions. **(A)** Comparison of the FeP:MoFeP docking geometry (DG) in ^t/o^Complex-1 (maroon) and ^t/o^Complex-2 (blue) observed in the DG2 configuration (black outline) compared to DG1 (gold; PDB ID: 2AFH) and DG3 (purple; PDB ID: 2AFI) configurations characterized by X-ray crystallography. **(B)** Structural overlay of ^t/o^Complex1 and ^t/o^Complex-2, indicating that the only large-scale conformational changes are observed in FeP. “Proximal” and “distal” refer to the *α*^1^*β* ^1^ and *α*^2^*β* ^2^ halves of MoFeP bound and not bound to FeP, respectively. The *α*III domains of MoFeP are outlined in gray. **(C)** Structural overlay of the FeP components in ^t/o^Complex1 and ^t/o^Complex-2, highlighting the nucleotide-dependent conformational differences (hinging - left, twisting - right) motions of the *γ*^1^ and *γ*^2^ subunits relative to one another during ATP hydrolysis. The axes of the *γ*100’s helices that radiate from [4Fe:4S] cluster are shown as black lines to illustrate these conformational differences. **(D**,**E)** Residues in the vicinity of FeMoco (**D**, *α*H274, *α*F300, *α*H451; **E**, *α*W253) that have undergone conformational changes in the distal subunit (*α*^2^) of MoFeP during turnover. Movements between the resting-state ^rs^MoFeP (gray) and the turnover structures (maroon and blue) are indicated with arrows.

The asymmetry present both in ATP hydrolysis and FePMoFeP interactions has important implications for the timing of ET events during catalysis. Compared to a concerted process, stepwise nucleotide hydrolysis by FeP would be expected to increase the lifetime of the activated DG2 complex and provide additional conformational states for orchestrating the multistep redox reactions that occur at FeMoco and the P-cluster^(6, 24)^. Similarly, an alternating docking mechanism between FeP and MoFeP (as imposed by negative cooperativity between the *αβ* halves) would effectively slow down successive ET steps to a given *αβ* subunit, providing sufficient time for anticipated protein and metallocluster rearrangements during N_2_ reduction. This is consistent with the suggestion of Thorneley and Lowe that the slow kinetics of nitrogenase (*k*_turnover_ ≈1 s^-1^) governed by FeP-MoFeP interactions may be a mechanistic imperative to favor N_2_ fixation over the competing but less demanding H_2_ reduction^(8)^.

Having established ^t/o^Complex-1 and ^t/o^Complex-2 as two distinct intermediates along the FeP cycle, we next investigated if they exhibited any conformational changes in their MoFeP components that may be correlated with the ATP hydrolysis state. In both ^t/o^Complex-1 and ^t/o^Complex-2, the P-clusters are in their fully reduced, all-ferrous (P^N^) forms, and the protein backbone arrangements of MoFeP’s in both complexes are essentially indistinguishable from one another as well as from those in ^rs^MoFeP and previously determined MoFeP crystal structures (C*α* RMSD: 0.2 Å; **Fig. 2B; figs. S7, S8; table S4**). We observed no large-scale conformational changes that could account for mechanical coupling and negative cooperativity between symmetry-related FeP docking surfaces on MoFeP, implicating the involvement of a dynamic allosteric mechanism (e.g., FeP-induced changes in MoFeP conformational entropy) **(fig. S9)**^(25)^. Yet, a detailed inspection of the structures revealed that the FeP-free (i.e., “distal”) *α*^2^*β*^2^ half in both ^t/o^Complex-1 and ^t/o^Complex-2 possessed several features that distinguish it from the FeP-bound (i.e., “proximal”) *α*^1^*β* ^1^ half and ^rs^MoFeP.

First, there are several, highly conserved residues (*α*^2^Trp253, *α*^2^His274, *α*^2^Phe300, *α*^2^His451)^(26)^ in the vicinity of the distal FeMoco which adopt non-resting-state conformations **(Figs. 2D,E)**. *α*Trp253 is particular in its cis-peptide bond to *α*Ser254 and its position in a proposed substrate access channel from the protein surface to FeMoco^(27, 28)^. The observed conformational flip in *α*^2^Trp253 leads to the diversion of this putative channel to an alternate face of FeMoco **(Fig. 2E, fig. S10)**. *α*^2^His274, *α*^2^Phe300 and *α*^2^His451 sidechains appear to have undergone a concerted motion compared to their resting state **(Fig. 2D)**, whereby *α*^2^His274 and *α*^2^Phe300 assume a similar configuration as that seen in the low-pH crystal structure of MoFeP **(fig. S11)**^(29)^. This *α*His274 configuration was proposed to form a water-bridged H-bond to a protonated belt sulfur (S5A) of FeMoco^(29)^. Along these lines, the observed rearrangement of the *α*^2^His274 sidechain in the turnover complexes could be envisioned to stabilize a protonated FeMoco intermediate and/or to increase the reduction potential of the cofactor, thus promoting its reduction by the P-cluster.

Second, the cryoEM densities surrounding the *α*^2^His442 and homocitrate ligands to the Mo center of the distal FeMoco are considerably less well defined compared to their counterparts in the proximal *αβ* half and the residues in the vicinity **(fig. S12)** and cannot be modeled with the resting-state configurations of these ligands **(Fig. 3)**. The reduction in map density is particularly pronounced for ^t/o^Complex-2 (i.e., midATP-hydrolysis) compared to ^t/o^Complex-1 (i.e., pre-ATP-hydrolysis). These observations suggest that *α*^2^His442 and homocitrate are mobile during turnover and that Mo undergoes changes in inner-sphere coordination in a way that is correlated with the nucleotide hydrolysis state of FeP bound to the opposing *αβ* half of MoFeP. The substitution of Mo with V or Fe in alternative nitrogenases, the replacement of homocitrate with citrate, and alterations in H-bonding to homocitrate have been shown to substantially diminish N_2_ reduction activity and alter substrate specificity^(2, 30-32)^. Indeed, the direct involvement of the Mo center in N_2_ reduction has been proposed early on^(2, 33)^, although recent experimental findings have shifted the focus to the central Fe centers of FeMoco as being the primary sites for substrate activation^(15, 16)^. Our cryoEM observations provide strong evidence that the Mohomocitrate moiety also directly participates in dynamic structural transformations that accompany catalysis at FeMoco.

**Figure 3.**
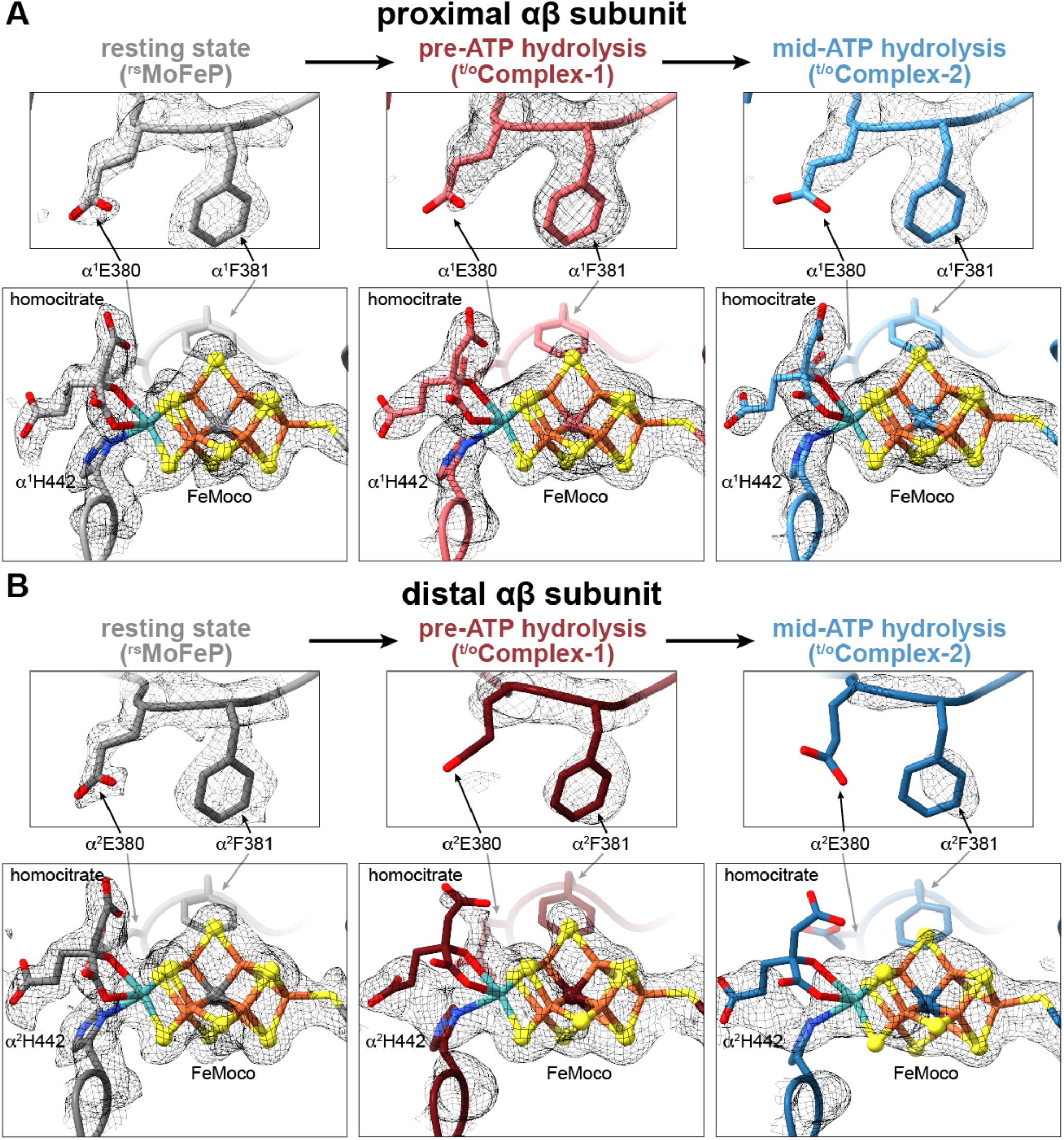
Changes in the FeMoco environment observed during catalytic turnover. **(A, B)** Views of FeMoco and the nearby residues *α*E380 and *α*F381 in the proximal **(A)** and distal **(B)** *αβ* halves of MoFeP in ^rs^MoFeP (gray), ^t/o^Complex-1 (maroon), and ^t/o^Complex-2 (blue) structures. CryoEM maps for each individual structure are contoured at the same level.

Third, portions of a large domain in the distal *αβ* subunit comprising residues *α*25-48 and *α*2378-403 (particularly in ^t/o^Complex-2) possess increased mobility compared to the rest of the MoFeP **(fig. S13)**. This so-called *α*III domain forms a lid above FeMoco and includes residues *α*Glu380 and *α*Phe381 that form close contacts with FeMoco **(Figs. 2B, 3)**. In resting state MoFeP, *α*III is well-ordered, *α*Glu380 forms water-bridged H-bonds to the Mo-ligated *α*His442 sidechain and homocitrate, and *α*Phe381 is in van der Waals contact with the labile belt sulfide S2B of FeMoco **(Fig. 3A)**. By contrast, in the distal *αβ* halves of ^t/o^Complex-1 and ^t/o^Complex2, the cryoEM densities for *α*III and sidechains of *α*^2^Glu380 and *α*^2^Phe381 sidechains are either diffuse or entirely missing, consistent with their movement during turnover **(Fig. 3B, fig. S13)**. This movement is likely coupled to the dynamics of the Mo ligands and FeMoco as a whole. *α*III has also been shown to undergo major structural rearrangements associated with the insertion of FeMoco into MoFeP^(34)^. Furthermore, *α*III displays some of the highest temperature factors in MoFeP crystal structures and is positioned away from lattice contacts **(table S5)**^(10, 11, 24, 29, 35)^, implying that it is inherently more flexible than other parts of MoFeP. Combined with this observation, ^t/o^Complex-1 and ^t/o^Complex-2 structures point to a possible role of *α*III mobility in nitrogenase catalysis. Notably, *α*III abuts the docking surface of MoFeP for MgADP-bound FeP in DG3 and may lie in the trajectory of FeP moving directionally across the MoFeP surface during ATP hydrolysis^(11)^. Thus, a dynamic *α*III domain could also provide a direct mechanical conduit between FeP and the proximal FeMoco, further linking the timing of nucleotide-dependent FeP-MoFeP interactions to redox transformations at FeMoco.

In conclusion, the cryoEM structures of FeP-MoFeP complexes formed during catalysis reveal that MoFeP is highly dynamic, opposing the existing view of its structural invariance derived from crystallographic studies. From the hydrolysis of ATP molecules to the multimodal FeP and MoFeP interactions, asymmetry and directionality emerge as pervasive elements in nitrogenase function, and may be critical for the timing of successive electron and proton transfers to FeMoco to optimize N_2_ reduction and avoid unproductive H_2_ evolution. Importantly, under turnover conditions, MoFeP displays several unforeseen structural features such as conformational changes in specific residues near FeMoco **(Figs. 2D, E)**, mobility in Mo ligands **(Fig. 3B)** and increased *α*III domain motions **(fig. S13)**, which are correlated with the nucleotide-state of a distally-bound FeP and can reasonably be linked to redox gating and catalytic events. Based on the available structures, it is not obvious how FeP-MoFeP interactions induce these conformational changes over a distance of 80 Å or which states of the MoFeP cycle (i.e., E_0_-E_8_) the observed nitrogenase complexes correspond to. It is safe to assume, however, that all copies of FeMoco in these complexes represent the more stable or longer-lived of the many catalytic intermediates present in the turnover solution. The density maps for the proximal FeMoco’s are essentially identical to those for the resting-state cofactor **(Fig. 3)**. Thus, an assignment of E_0_ for the proximal cofactors is plausible, although E1-E4 are also reasonable as these states have been proposed as hydride-bound FeMoco intermediates^(15, 16)^, which may be structurally indistinguishable from E_0_ at the current resolution of these cryoEM maps. At the same time, the obvious deviations between the densities of distal and proximal FeMoco’s suggest that the distal cofactors in ^t/o^Complex-1 and ^t/o^Complex-2 represent an Ex state or a mixture of Ex states that are different and likely more advanced in the catalytic cycle (i.e., E1) than the proximal FeMoco’s, and involve the participation of the Mo center. In light of the asymmetry between the FeMoco’s in the two *αβ* halves, it is tempting to propose a “ping-pong”-type mechanism in which the cofactors proceed through each of the eight catalytic steps in an alternating fashion. This scenario would assign a dual role to FeP: (1) to deliver an electron to one *αβ* subunit of MoFeP and (2) to suppress FeP binding to the opposite *αβ* subunit while priming it for catalytic transformations through long-distance activation of electron, H_2_ and/or substrate access pathways to the distal FeMoco. Although the examination of such a mechanism will require future studies, our current work illustrates that it is finally possible to characterize *bona fide* intermediates of nitrogenase catalysis at near-atomic resolution via cryoEM, representing a critical step toward understanding the mechanism of this enigmatic enzyme in full structural detail.

## Acknowledgements

We thank K. Corbett, S. Narehood, J. Figueroa, and R. Subramanian for critical discussions, and members of the Tezcan and Herzik Labs for their assistance. Molecular graphics and analyses were performed with UCSF ChimeraX, developed by the Resource for Biocomputing, Visualization, and Informatics at the University of California, San Francisco, with support from National Institutes of Health grant R01-GM129325 and the Office of Cyber Infrastructure and Computational Biology, National Institute of Allergy and Infectious Diseases. We also think members of UCSD’s CryoEM facility, the Stanford-Slac Cryo-EM Center (S^2^C^2^), and UCSD’s Physics Computing Facility for help in data collection, data processing, and computational support.

## Author contributions

Conceptualization: HLR, MAH, FAT

Methodology: HLR, BDC, MAH, FAT

Investigation: HLR, BDC, HPMN, MAH

Visualization: HLR, BDC, MAH

Funding acquisition: MAH, FAT

Project administration: MAH, FAT

Supervision: MAH, FAT

Writing - original draft: HLR, MAH, FAT

Writing - review & editing: HLR, BDC, MAH, FAT

## Data and materials availability

Structural models have been deposited in the Protein Data Bank (PDB) with accession codes 7UT6 (^rs^MoFeP, *C*_*1*_ symmetry), 7UT7 (^rs^MoFeP, *C*_*2*_ symmetry), 7UT8 (^t/o^Complex-1), 7UT9 (^t/o^Complex-2), and 7UTA (BeF_x_-trapped complex). The corresponding cryoEM maps are available at the Electron Microscopy Data Bank (www.ebi.ac.uk/emdb/) with accession codes EMD-26756 (^rs^MoFeP, *C*_*1*_ symmetry), EMD-26757 (^rs^MoFeP, *C*_*2*_ symmetry), EMD-26758 (^t/o^Complex-1 consensus MoFeP), EMD-26759 (^t/o^Complex-1 locally refined FeP), EMD-26760 (^t/o^Complex-1), EMD-26761 (^t/o^Complex-2 consensus MoFeP), EMD-26762 (^t/o^Complex-1 locally refined FeP), EMD-26763 (^t/o^Complex-2), and EMD-26764 (BeF_x_-trapped complex). All other data are available in the main text or the supplementary materials.

