## Supplemental Materials for "CryoEM Structures of the Nitrogenase Complex During Catalytic Turnover"

### Supplementary Materials

Materials and Methods
Figs. S1 to S13
Tables S1 to S5
References (1 - 14)

#### Materials and methods

##### Protein expression and purification

Wild-type, untagged FeP and MoFeP were expressed in their native organism, *Azotobacter vinelandii* (*Av*) cells (strain DJ) using previously established protocols[(1)]. Briefly, Av cultures were grown aerobically in Burk's media (181 mM sucrose, 0.9 mM $CaCl_2$, 1.7 mM $MgSO_4$, 35 $\mu$M $FeSO_4$, 2 $\mu$M $Na_2Mo_2O_4$, 0.2 mM citric acid, 10 mM $K_3PO_4$ pH 7.5, 3 mM $NH_4Cl$) in a 60 L fermenter at 30 C, 200 rpm. Cells were harvested and pelleted ~4 h after derepression of nitrogenase, as indicated by a spike in dissolved oxygen content. Cell pellets were stored at -80 C until purification.

Cell lysis and protein purification were carried out under ultrahigh-purity Ar on a Schlenk line, or under a 95% Ar/5% $H_2$ mixture in a Coy Lab anaerobic chamber using previously established protocols[(1)]. All buffers used were purged of air and stored under Ar. Cell pellets were resuspended in ~200 mL equilibration buffer (50 mM Tris pH 7.75, 200 mM NaCl, 5 mM sodium dithionite (NaDT), 0.1 mg/mL DNase I) prior to lysis with a microfluidizer at 16,000 psi Ar. The lysate was centrifuged at 12,000 rpm for 75 min. Both nitrogenase component proteins were purified from the supernatant by separation on a DEAE Sepharose column with a NaCl gradient (200 to 500 mM NaCl in 50 mM Tris pH 7.75, 5 mM NaDT). MoFeP eluted at ~25 mS/cm, and FeP eluted at ~30 mS/cm. Fractions containing nitrogenase proteins were brown in color and were verified by sodium dodecyl sulfate polyacrylamide gel electrophoresis (SDS-PAGE). Fractions containing FeP or MoFeP proteins were pooled and diluted two-fold with salt-free buffer (50 mM Tris, pH 7.75), then concentrated using a second, smaller DEAE Sepharose column by eluting with high salt buffer (500 mM NaCl, 50 mM Tris, pH 7.75, 5 mM NaDT). MoFeP and FeP were further purified with a Sepharose 200 gel filtration column (500 mM NaCl, 50 mM Tris, pH 8.0, 5 mM NaDT). Fractions containing pure protein were identified with SDS-PAGE. Purified protein was concentrated using an Amicon concentrator at 20 psi 95% Ar/5% $H_2$ using a 30 kDa and 100 kDa cutoff membrane for FeP and MoFeP, respectively. Purified proteins were syringe filtered through a 0.2 µm filter membrane. Protein concentrations were determined using Bradford assay and verified with an Fe chelation assay (6.2 M guanidine-HCl, 2 mM 2,2'-bipyridine, 10% glacial acetic acid) by measuring absorption at 522 nm and using an extinction coefficient of 8650 $M^{-1}$ $cm^{-1}$, using 4 Fe per FeP and 30 Fe per MoFeP in stoichiometric calculations. Purified proteins were determined to be fully active for $C_2H_2$ and $N_2$ reduction assays that were performed as previously described[1]. Purified proteins were stored under liquid $N_2$ and underwent only one freeze-thaw cycle before use.

##### Sample preparation for EM analysis

All EM samples were prepared under ultra-high-purity $N_2$. FeP and MoFeP were buffer exchanged into reaction buffer (20 mM Tris, pH 8.0, 25 mM NaCl, anaerobic) and concentrated using 30 kDa and 100 kDa Microcon centrifugal filters, respectively. Protein concentrations were measured using an Fe chelation assay. All EM samples contained a final concentration of 6 µM MoFeP, 60 µM FeP (except for the MoFeP control sample, which did not contain FeP), 5 mM $MgCl_2$, 5 mM $Na_2$ATP, 5 mM NaDT, 20 mM Tris, pH 8.0, and 25 mM NaCl. All reaction component stock solutions were made, degassed, and syringe filtered (0.2 $\mu$m filter) immediately prior to use. In addition, the $BeF_x$-inhibited sample contained 25 mM NaF and 5 mM $BeSO_4$. Proteins were transferred into sealed reaction vials using Hamilton gas-tight syringes after reaction components had been mixed. Catalysis was initiated in the turnover and $BeF_x$-inhibited sample by addition of FeP after all other components had been mixed. 10 $\mu$L of each EM sample was transferred to a 200 $\mu$L thin-wall tube under nitrogen and immediately flash frozen in liquid nitrogen 15 seconds after reaction was initiated (or after last component was added in the case of MoFeP control). Flash frozen EM samples were stored under liquid $N_2$ until grid preparation. All samples were prepared on UltraAuFoil 1.2/1.3, 300 mesh grids that had been freshly plasma-cleaned using a Gatan Solarus II plasma cleaner (10 s, 15 Watts, 75% Ar/25% $O_2$ atmosphere). To minimize exposure to air and reaction time between thawing frozen samples and freezing grids, all grids were prepared using a custom manual plunge freezer designed by the Herzik Lab located in a humidified (>95% relative humidity) cold room (4 C). Immediately after thawing, 3 $\mu$L of the sample was applied to the grid surface followed by manual blotting for ~5 to 6 s using Whatman No. 1 filter paper before vitrifying in an 50% ethane/ 50% propane liquid mixture cooled by liquid $N_2$[2]. The time that each sample spent outside of liquid $N_2$ was less than 15 sec. Grids were stored under liquid $N_2$ until data collection.

##### EM data acquisition and data processing

**$^{rs}$MoFeP:** Data acquisition for the free MoFeP ($^{rs}$MoFeP) was carried out at UCSD's CryoEM Facility on a Titan Krios G3 (Thermo Fisher Scientific) operating at 300 keV equipped with a Gatan BioContinuum energy filter. Images were collected at a magnification of 165,000x in EF-TEM mode (0.815 Å calibrated pixel size) on a Gatan K2 detector using a 20-eV slit width and a cumulative electron exposure of ∼65 electrons/Å$^2$ (50 frames). Data were collected automatically using EPU with aberration free image shift using a defocus range of -0.5 - -2.5 $\mu$m. Motion correction was performed using the MotionCor2 frame alignment program implemented within RELION(3) 4.0-beta1 using 7x7 tiled frames with a B-factor of 250. Dose-weighted images were used for preliminary processing and CTF estimation using CTFFind4 within RELION (1024-pixel box size, 0.1 amplitude contrast, 30 Å minimum resolution, 3 Å maximum resolution)[(3,4)]. Aligned images with a CTF-estimated resolution below 5 Å or with a cumulative total motion exceeding 60 Å were excluded. For free MoFeP, initial particle picks were obtained using cryoSPARC Live's blob picker (50 - 120 Å circular and elliptical blobs) and an *ab initio* model was generated using optimal 2-D classes[(5)]. This initial model was then used to generate 2-D templates for automated template-based particle picking using RELION 4.0-beta2[(3)]. A total of (1,527,385+1,508,038) particle picks were extracted from (1,587+2,337) micrographs collected across two different sessions from two different grids, downsampled 4 x 4 (3.26 Åpixel, 64 pixel box size) and subjected to iterative rounds of reference-free 2-D classification (100 classes, tau_fudge=1, VDAM, ignore first CTF peak, 140 Å mask). Particles were subjected to four iterative rounds of 2-D classification and those 2-D class averages containing the strongest secondary structural details were isolated (1,497,616 particles in total) for 3-D auto-refinement using $C_1$ symmetry[(6)]. Each session was then processed in parallel. The refined coordinates were used to re-center and re-extract particles unbinned (0.815 Å/pixel, 384 pixel box size). These particles were refined against a scaled version of the previously refined map followed by CTF refinement (per-particle defocus UV, per-micrograph astigmatism, anti-symmetrical and symmetrical higher-order aberrations). Following an iterative rounds of 3-D and CTF refinement, particles were subjected to RELION's Bayesian particle polishing using parameters trained against the data (–s_vel 1.52100 –s_div 15030.00000 –s_acc 2.35500)[(3)]. Following particle polishing, 3-D auto-refinement, and CTF-refinement, a 2.12 Å structure was obtained. These particles were then subjected to a no-alignment 3-D classification (8 classes, tau_fudge=2) and the best classes (167,110 and 214,991 particles) were selected for iterative rounds of 3-D and CTF refinement followed by particle polishing using the same parameters but 512-pixel extraction box size. Both sessions were then combined and a 3-D auto-refinement led to a to ∼2.01 Å refinement. Another round of no-alignment 3-D classification was performed (6 classes, tau_fudge=8) and particles comprising the highest-quality classes (177,123 particles) were combined 3-D auto-refined and then imported into cryoSPARC for a non-uniform refinement using $C_1$ symmetry (1.91 Å resolution) or $C_2$ symmetry (1.81 Å resolution)[(5)].

**$^{t/o}$Complex-1 and $^{t/o}$Complex-2:** Data for the nitrogenase turnover sample were collected at the S$^2$C$^2$ Stanford-SLAC CryoEM Center on TEM Gamma (Titan Krios G3i (Thermo Fisher Scientific) equipped with a Gatan K3 direct electron detector) operating at 300 keV. Images were collected at a magnification of 135,000x (0.835 Å/pixel) on a K3 detector with an electron exposure of ∼65 electrons/Å$^2$ (66 frames) with a nominal defocus range of -1.2 - -2.0 $\mu$m. Motion correction was performed using the MotionCor2 frame alignment program implemented within RELION 4.0-beta1 using 10x14 tiled frames with a B-factor of 250[(3,4)]. Dose-weighted images were used for preliminary processing and CTF estimation using CTFFind4 within RELION (1024-pixel box size, 0.1 amplitude contrast, 30 Å minimum resolution, 3 Å maximum resolution)[(3,7)]. Aligned images with a CTF-estimated resolution below 5 Å or with a cumulative total motion exceeding 60 Å were excluded. The resting state MoFeP structure was used to template pick ∼50 movies and the top picks were used to train crYOLO for picking against the entire data set[(8)]. 19,711,170 picks were obtained from 14,903 micrographs and extracted in RELION 4.0-beta2 downsampled 8 x 8 (6.68 Å/pixel, 64 pixel box size), randomly split into ∼1M particle sets, and each subjected to iterative rounds of reference-free 2-D classification (200 classes, tau_fudge=1, VDAM, ignore first CTF peak, 180 Å mask) where only obvious false classes were eliminated[(3)]. 11,955,963 particles were then re-centered and re-extracted, downsampled 4 x 4 (3.34 Å/pixel, 96 pixel box size), randomly split into ∼1M particle sets and each subjected to iterative rounds of reference-free 2-D classification (200 classes, tau_fudge=1, VDAM, ignore first CTF peak, 180 Å mask) where only obvious false classes were eliminated. 7,708,206 particles from the best classes were combined, randomly split into 10 subsets, and subjected to another round of 2-D classification. The best nitrogenase classes were then set aside and the remaining classes were re-ran through 2-D classification. The best nitrogenase classes were then combined with the previous run and subjected to 3-D auto-refine. These particles were then re-centered and re-extracted downsampled 4 x 4 (3.34 Å/pixel, 96 pixel box size) with duplicates removed. 4,121,671 particles were imported into cryoSPARC v3.3.2 and subjected to a heterogeneous refinement using four nitrogenase 1:1 classes and one 20S proteasome class (EMDB-8741)[(5)]. The best 1:1 nitrogenase class comprising 2,511,497 particles was then subjected to a 2-class heterogenous refinement using 1:1 nitrogenase and MoFeP volumes as initial models. 1:1 complexes and MoFeP particles were then re-ran through this 2-class heterogenous refinement two times before combining all the 1:1 complexes and MoFeP particles separately and subjected to a non-uniform Refinement. These particles were then re-centered and re-extracted in RELION downsampled 2 x 2 (1.67 Å/pixel, 192 pixel box size) with duplicates removed[(3)]. 906,326 1:1 complex particles were subjected to a non-uniform refinement in cryoSPARC yielding a Nyquist-limited 3.43 Å resolution map with high-quality FeP density. These particles were then subjected to a 3-D variability analysis (two modes, four intermediate clusters, 5 Å low-pass, no overlap)[(9)]. The best 1:1 nitrogenase class was then subjected to another round of non-uniform refinement and 3-D variability analysis (two modes, four intermediate clusters, 5 Å low-pass filter)(9). Each cluster was then independently subjected to non-uniform refinement and the best two classes with FeP density for both subunits were re-centered, re-extracted in RELION without downsampling (0.835 Å/pixel, 384 pixel box size) and 3-D auto-refined followed by Bayesian particle polishing using parameters trained against the data (–s_vel 0.9225 –s_div 6570.00000 –s_acc 2.65500)(3). These particles were then imported into cryoSPARC for a non-uniform refinement, yielding 2.38 Å and 2.34 Å resolution maps for the ATP-ATP and ATP/ADP-ADP structures, respectively(5). A soft mask for FeP was then used for a local refinement (4-Å deviation over priors, 4- search, 4-Å shift search) yielding 2.75 Å and 3.01 Å maps for the ATP-ATP and ATP/ADP-ADP structures, respectively. The composite half maps from each independent half set from full and locally refined were assembled (maximum voxel value) and subjected to deepEMhancer(10) (high-resolution model; version 0.13). The FSC estimated resolution for the composite maps 2.28 Å and 2.29 Å ATP-ATP and ATP/ADP-ADP structures, respectively.

**$BeF_x$-trapped nitrogenase complex:** Data for the $BeF_x$-trapped complex were collected at UCSD's CryoEM Facility on a Titan Krios G3 (Thermo Fisher Scientific) operating at 300 keV equipped with a Gatan BioContinuum energy filter. Images were collected at a magnification of 165,000x in EF-TEM mode (0.815 Å calibrated pixel size) on a Gatan K2 detector using a 20-eV slit width and a cumulative electron exposure of ~65 electrons/Å$^2$ (50 frames). Data were collected automatically using EPU with aberration free image shift using a defocus range of -0.5 - -2.5 $\mu$m. 4 separate data sets were collected using 0, 15 or 25 specimen tilt. Motion correction was performed using the MotionCor2 frame alignment program implemented within RELION 4.0-beta1 using 7x7 tiled frames with a B-factor of 250(4). Dose-weighted images were used for preliminary processing and CTF estimation using CTFFind4 within RELION (1024-pixel box size, 0.1 amplitude contrast, 30 Å minimum resolution, 3 Å maximum resolution)(3). Aligned images with a CTF-estimated resolution below 5 Å or with a cumulative total motion exceeding 60 Å were excluded. Initial particle picks were obtained using RELION's template picker using free MoFeP as a template(3). A total of (271,261+165,767+349+541+330,486) particle picks were extracted from (2,085+1,718+2,211+2,099) micrographs collected across four different sessions from two different grids, downsampled 8 x 8 (6.52 Å/pixel, 48 pixel box size). Each data set was subjected to two rounds of reference-free 2-D classification (50 classes, tau_fudge=1, VDAM, ignore first CTF peak, 220 Å mask). Particles from 2-D class averages containing the strongest secondary structural details (148,743+95,336+163,920+91,161 particles) were combined and subjected to another round of 2-D classification (50 classes, tau_fudge=1, VDAM, ignore first CTF peak, 220 Å mask). 424,249 particles were 3-D auto-refined ($C_1$ symmetry), re-centered and re-extracted (removing duplicates) without downsampling (0.815 Å/pixel, 384 pixel box size). The particles were then 3-D auto-refined and subjected to Bayesian particle polishing using parameters determined from the free MoFeP data set. These particles then underwent 3-D auto-refinement, CTF refinement (defocus UVA and aberrations), and a 3-D auto-refinement before a no-alignment 3-D classification (8 classes, tau_fudge=8). The best classes (397,392 particles) were then refined followed by local FeP masked 3-D classification (8 classes, tau_fudge=2). 2:1 complex particles were separated and 3-D refined followed by a subsequent no-alignment 3-D classification (4 classes, tau_fudge=24). The three best classes, representing 72,125 particles, were then subjected to 3-D autorefinement, CTF refinement, and a 3-D auto-refinement particles were subjected to Bayesian particle polishing using parameters determined from this data set (–s_vel 1.3275 –s_div 5955.00000 –s_acc 1.63500). A final series of 3-D auto-refinement, CTF refinement, and 3-D auto-refinement yielded a 2.40 Å resolution structure for the 2:1 $BeF_x$-trapped FeP:MoFeP complex.

Local resolution estimates were performed using cryoSPARC(5). 3-D FSC calculations were performed using the 3-DFSC server. Visualization was performed using UCSF's Chimera and ChimeraX. Particle meta data manipulation was performed using csparc2star.py and in-house developed Python scripts.

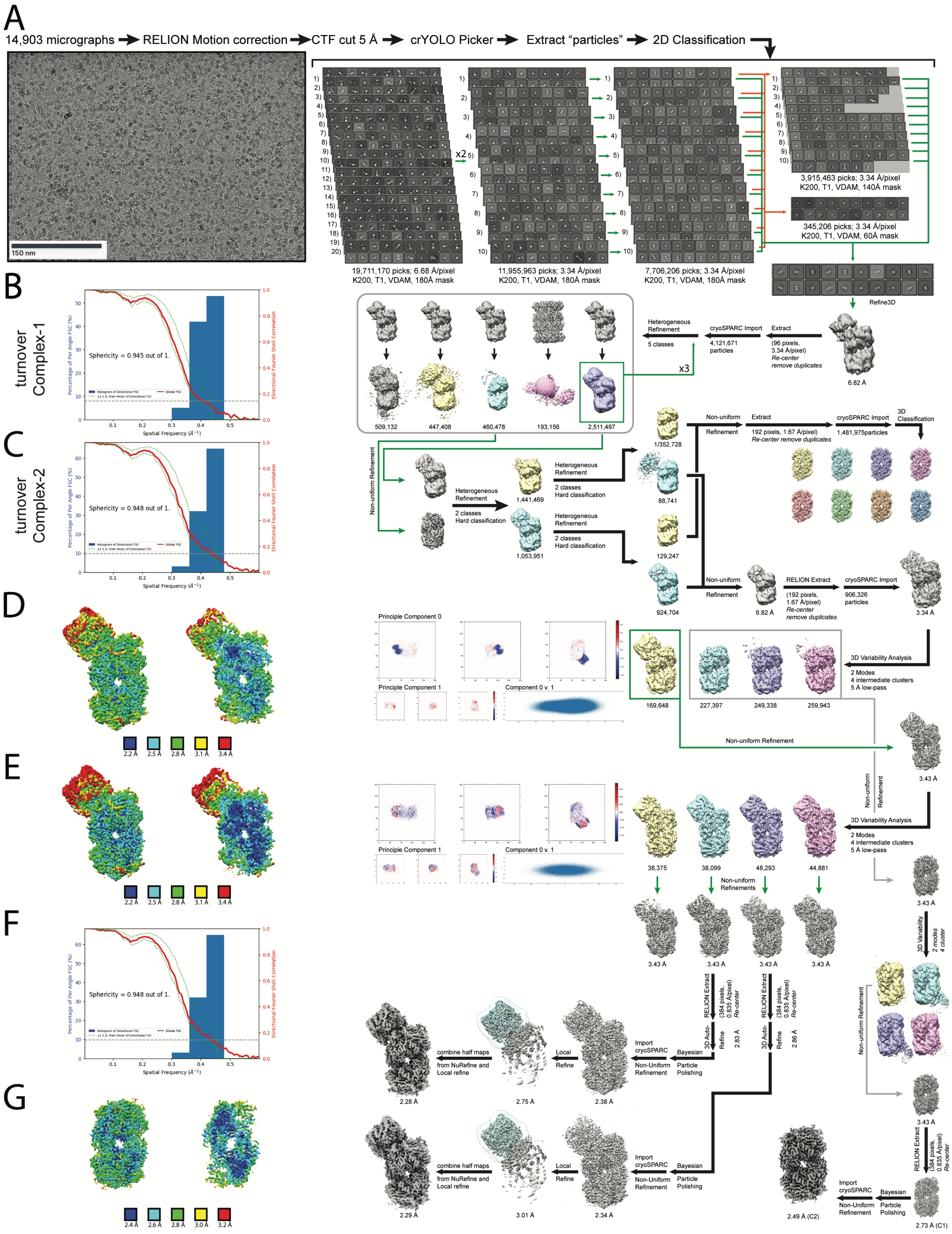


**Supplementary Figure S1. (caption on next page).**

**Figure S1. Data processing flowchart for the single-particle cryoEM analysis of nitrogenase complexes formed under turnover. (A)** Representative motion-corrected micrograph of vitrified nitrogenase collected at ∼1.5 µm underfocus. 19,711,170 particles were identified from dose-weighted micrographs by crYOLO trained using resting state MoFeP. These particles were extracted and downsampled 8 x 8 in RELION, randomly split into ∼1M particle sets, and subjected to iterative rounds of reference-free 2-D classification in RELION. Representative 2-D class averages are shown for each iterative step. The best nitrogenase classes were set aside (green arrows) while the remaining classes were randomly split into 10 subsets and subjected to another round of 2-D classification (orange arrows). The best classes were then combined and 3-D auto-refined to 6.82 Å. Particles were then imported into cryoSPARC and subjected to a 5-class heterogenous refinement. The best class for the 1:1 nitrogenase complex was selected (green box) and two iterative rounds of two class heterogenous refinements were performed. 1:1 FeP:MoFeP complex particles were isolated and re-extracted in RELION and imported into cryoSPARC for 3-D variability analysis. The cluster with the strongest FeP density in the 1:1 FeP:MoFeP complex was selected (green box) and non-uniform refined before a final round of 3-D variability analysis. The two clusters with density for both proteins were re-centered and re-extracted in RELION without downsampling, 3-D auto-refined, and subjected to Bayesian particle polishing. After polishing, the particles were imported into cryoSPARC and locally refined. The half maps were combined, resulting in the $^{t/o}$Complex-1 and $^{t/o}$Complex-2 resolving to 2.28 Å and 2.29 Å, respectively. Histogram and directional 3-D FSC plots generated from the independent composite half maps contributing to the ∼2.28 Å and 2.29 Å resolution **(B)** $^{t/o}$Complex-1 and **(C)** $^{t/o}$Complex-2 structures, respectively. **(D)** EM density of $^{t/o}$Complex-1 colored by local resolution. The left image corresponds to the surface of $^{t/o}$Complex-1, and the right image is a cross-section of the complex. **(E)** EM density of $^{t/o}$Complex-2 colored by local resolution. The left image corresponds to the surface of $^{t/o}$Complex-2, and the right image is a cross-section of the complex. **(F)** Histogram and directional 3-D FSC plots generated from the independent composite half maps contributing to the ∼2.59 Å resolution MoFeP ($C_2$ symmetry) map. **(G)** EM density of MoFeP (C2 symmetry) colored by local resolution. The left image is surface view of MoFeP, and the right image is a cross-section of the protein.

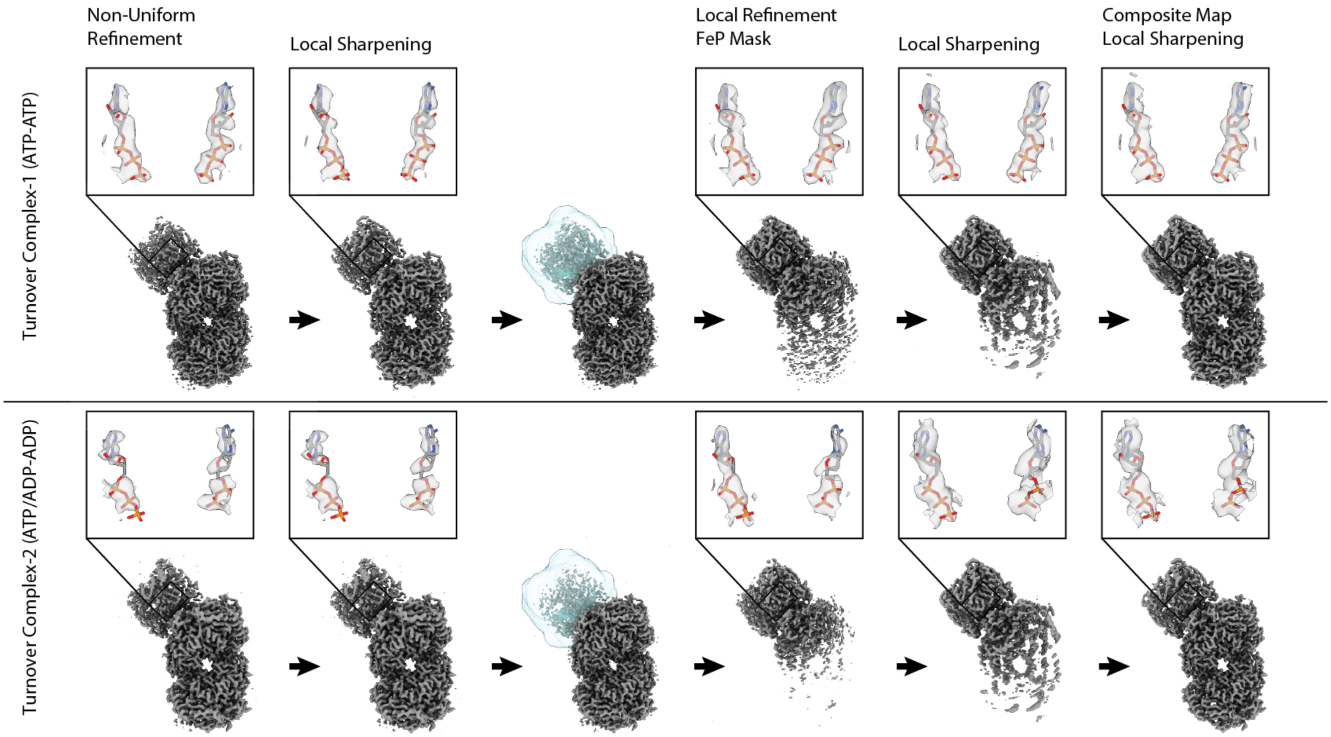


**Figure S2. Improvements in map quality for the 1:1 FeP:MoFeP complexes formed under turnover.** (left to right) Non-uniform refinements of the $^{t/o}$Complex-1 (*top*) and $^{t/o}$Complex-2 (*bottom*) yielded ~2.4 Å resolution maps with lower resolution regions for the FeP subunits. Local noise estimates and sharpening using deepEMhancer were used to improve the quality of the FeP density. Local refinement of each particle set using a soft FeP protein mask and subsequent local sharpening improved the EM density quality for the FeP subunits with lower quality density for the MoFeP subunits. Maximal voxel values for the non-uniform and local refinements were taken to generate composite half maps that were then used for resolution estimation, local noise estimates and sharpening.

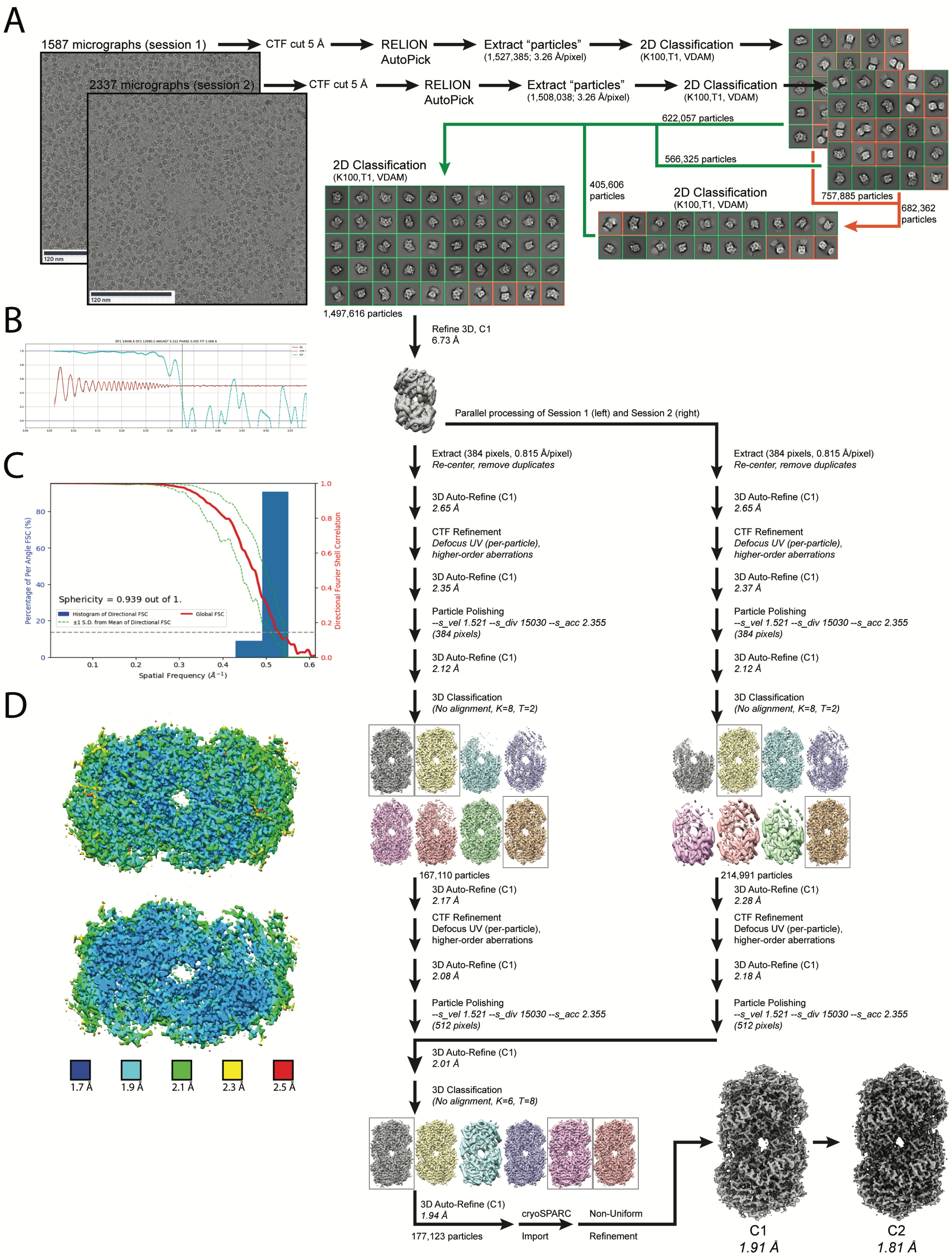


**Supplementary Figure S3. (caption on next page).**

**Figure S3. Data processing flowchart for the single-particle cryo-EM analysis of free MoFeP (rsMoFeP). (A)** Representative motion-corrected micrographs of vitrified MoFeP collected at ∼1.5 μm underfocus. ∼1.5M particles were identified using RELION's automated template based Autopicking, downsampled 8 x 8, and subjected to iterative rounds of reference free 2-D classification. Classes with strong secondary structural detail were isolated (green boxes/arrows), while poorly aligning classes were subjected to a final round of 2-D classification (orange boxes/arrows). All good classes were combined for an initial round of 3-D auto-refinement that refined to 6.73 Å resolution. Particles were then split into their respective sessions and processed in parallel. The refined coordinates were used to re-center and re-extract particles without binning and subjected to 3-D auto-refinement, CTF refinement, and 3-D auto-refinement before Bayesian particle polishing in RELION. No alignment 3-D classification was performed, and the best classes were selected (boxed). After iterative rounds of refinement, the sessions were combined, imported into cryoSPARC for non-uniform refinement in $C_1$ or $C_2$ symmetries using defocus and aberration refinement to yield 1.91 Å $C_1$ and 1.81 Å $C_2$ resolution structures. **(B)** Representative 2-D CTF fit for the data. **(C)** Histogram and directional 3-D FSC (11)plots generated from the independent half maps contributing to the ∼1.81 Å $C_2$ structure. **(D)** EM density of the $C_2$-refined structure colored by local resolution. The top image corresponds to the surface of MoFeP and the bottom image corresponds to a cross-section of the protein, highlighting core regions.

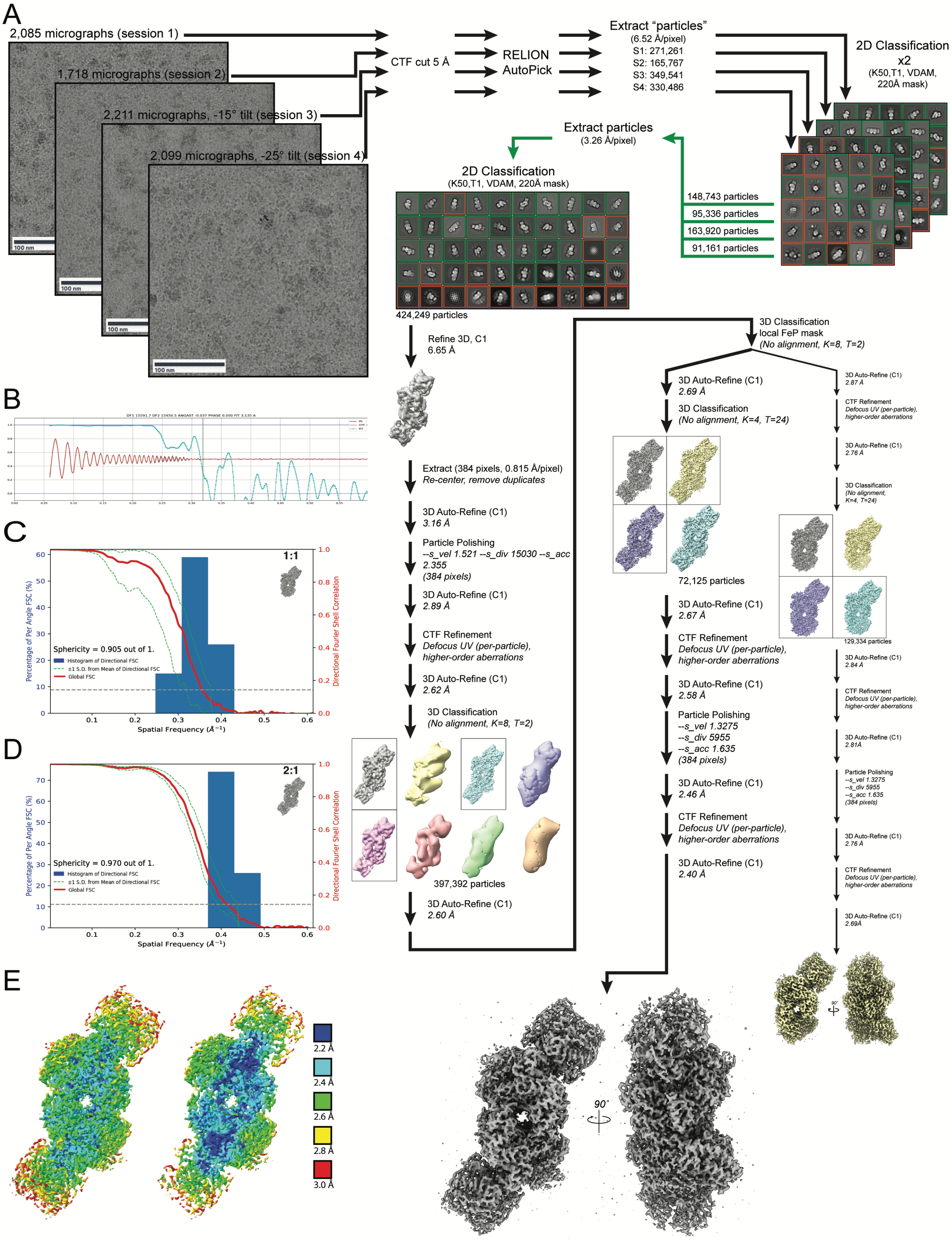


**Supplementary Figure S4. (caption on next page).**

**Figure S4. Data processing flowchart for the single-particle cryo-EM analysis of the $BeF_x$-trapped nitrogenase complex. (A)** Representative motion-corrected micrograph for each of the datasets for vitrified $BeF_x$-trapped nitrogenase complexes collected at ∼1.5 μm underfocus. A total of 1,117,055 particles identified from dose-weighted micrographs using RELION auto-pick with resting state MoFeP as a template. Particles were extracted from each dataset downsampled 4 x 4 and subjected to iterative rounds of 2-D classification in parallel before being combined for additional 2-D classification and 3-D auto-refinement. Particles were then re-centered and re-extracted unbinned and subjected to iterative rounds of 3-D auto- and CTF refinement, followed by a no alignment 3-D classification. The best classes were combined and subjected to additional rounds of no-alignment classification using a soft mask for the FeP subunits. 1:1 and 2:1 FeP:MoFeP complexes were separated and processed in parallel. After iterative rounds of 3-D auto- and CTF refinement, Bayesian particle polishing, ∼2.69 Å and ∼2.40 Å resolution structures were obtained for the 1:1 and 2:1 complexes, respectively. **(B)** Representative 2-D CTF fit for the data. (C-D) Histogram and directional 3-D FSC plots (11) generated from the independent half maps contributing to the structures of the **(C)** 1:1 and **(D)** 2:1 complexes **(E)** EM density for the 2:1 $BeF_x$-trapped FeP:MoFeP complex colored by local resolution. The left image shows the surface of the complex and the right image is a volume cross-section highlighting core regions.

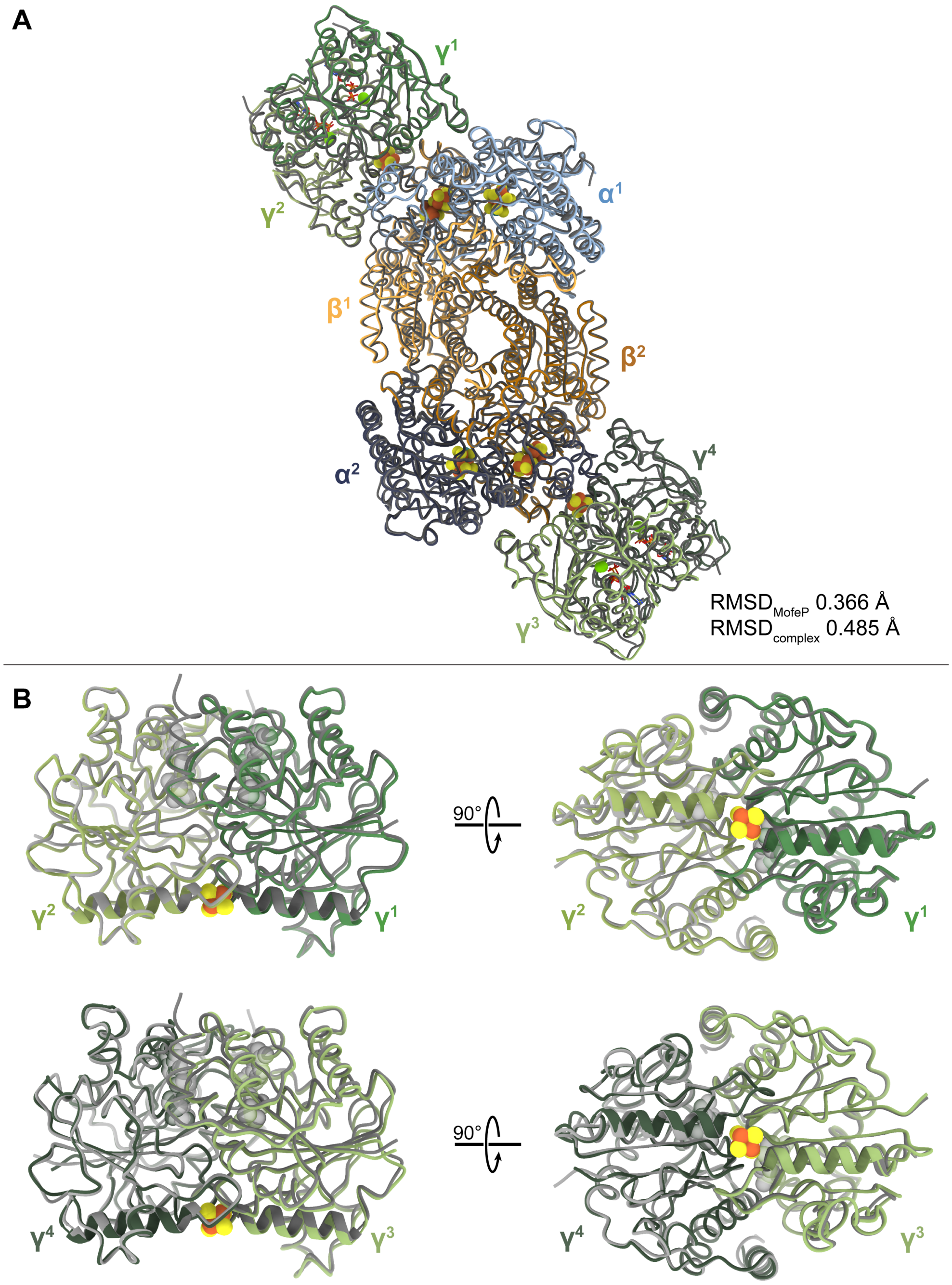


**Figure S5. Comparison of $BeF_X$-bound cryoEM structure with $AlF_X$ crystal structure.** **(A)** Overlay of $BeF_X$-trapped cryoEM structure (green, orange, and blue) with the $AlF_X$-trapped crystal structure (PDB ID: 1M34) highlighting their overall similarities. Root-mean-square-deviations (RMSDs) based on all C$\alpha$ positions in MoFeP and the entire complex are 0.366 Å and 0.485 Å, respectively. **(B)** Structural overlay of the FeP components ($\gamma^1$ and $\gamma^2$ top, ($\gamma^3$ and $\gamma^4$ bottom) in $BeF_X$- and $AlF_X$- trapped complexes, in which the left subunit ($\gamma^2$ or $\gamma^4$) has been aligned. The $\gamma$100's helices are depicted as ribbons.

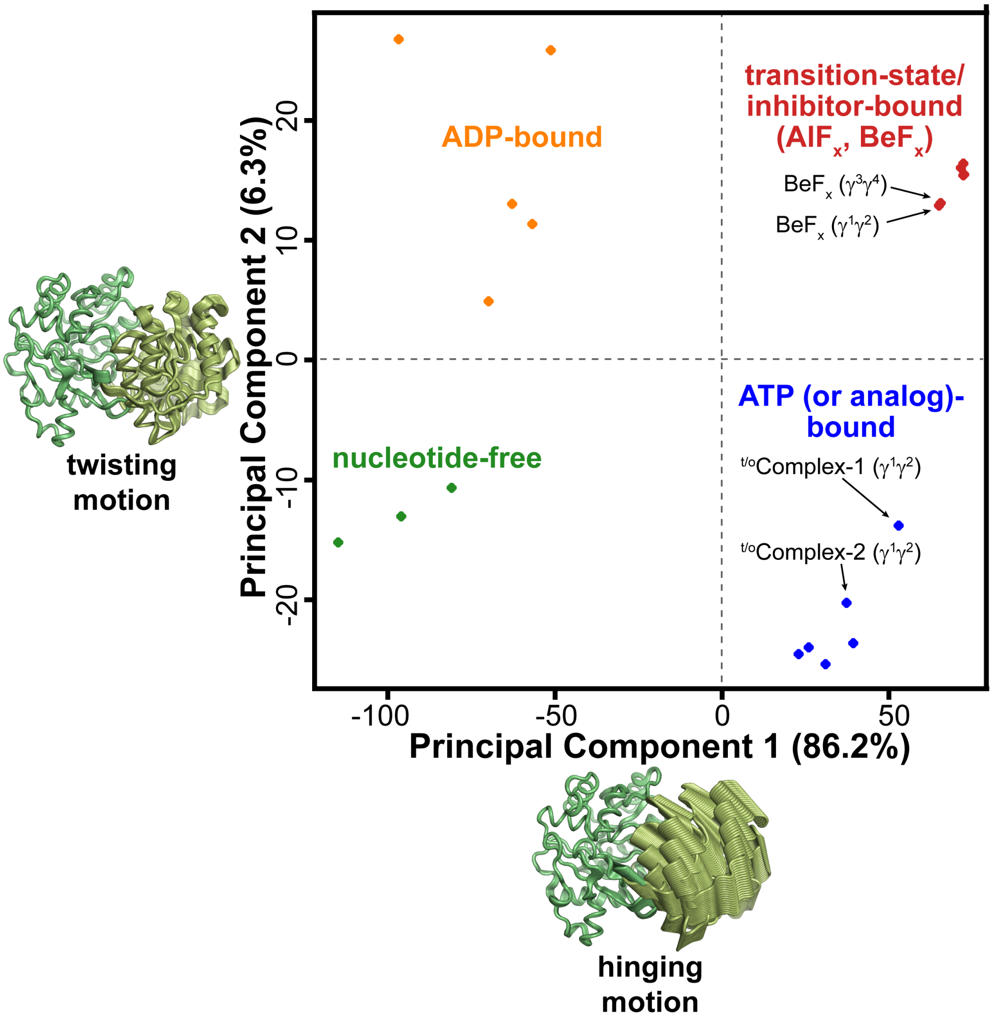


**Figure S6. Principal component analysis (PCA) of FeP.** PCA of FeP was carried out using twenty available free- or MoFeP-complexed FeP structures based on prior X-ray crystallographic or current cryoEM characterization. The first two principal components (PC) account for 92.5% of the variance. PC1 and PC2 are described by the hinging/rotation and twisting motions of the two $\gamma$-subunits with respect to one another. FeP conformations clustered into four nucleotide-state-dependent classes. FeP structures from the cryoEM structures indicated with arrows.

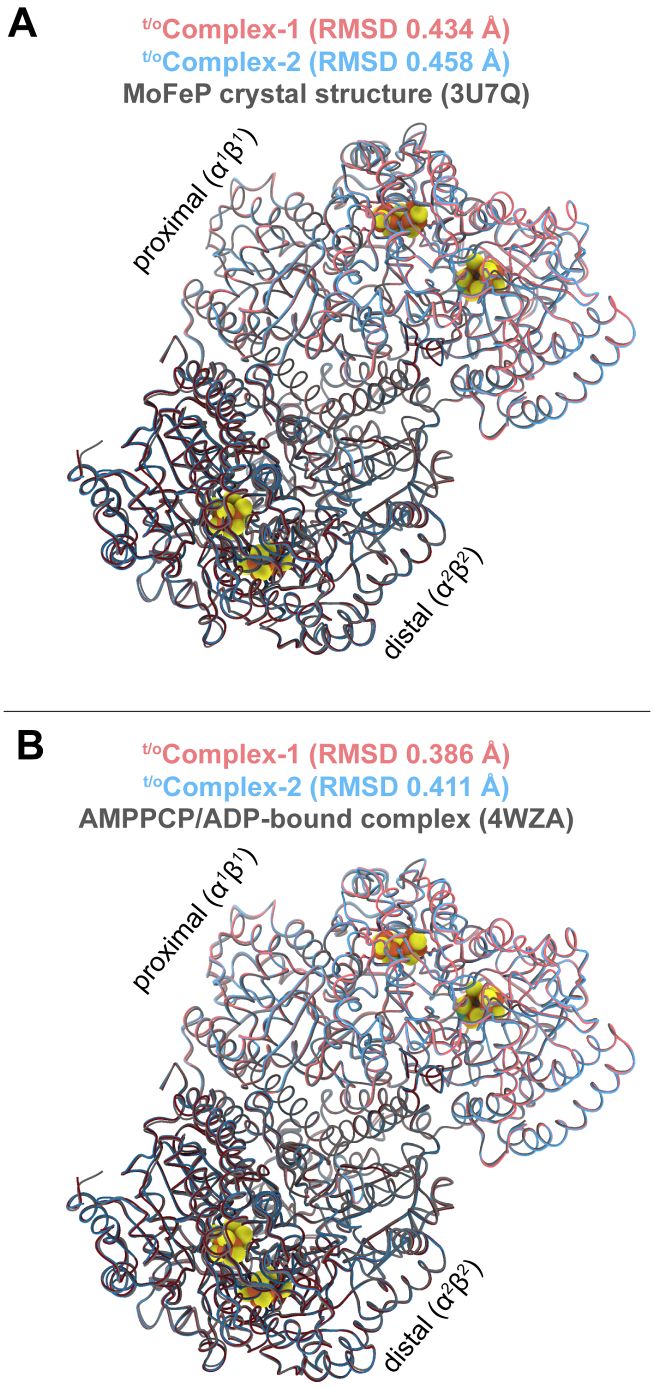


**Figure S7. Structural overlay of the MoFeP components of turnover complexes with MoFeP from previously determined crystal structures.** Overlay of $^{t/o}$Complex-1 (maroon) and $^{t/o}$Complex-2 (blue) with **(A)** a crystal structure of MoFeP (gray, PDB ID: 3U7Q), and **(B)** MoFeP from the AMPPCP/ADP bound FeP-MoFeP complex crystal structure (gray, PDB ID: 4WZA chains A,B,C,D). Root-mean-square-deviations (RMSDs; based on all C$\alpha$ positions) between the MoFeP components of the turnover complexes and those from the crystal structures are indicated.

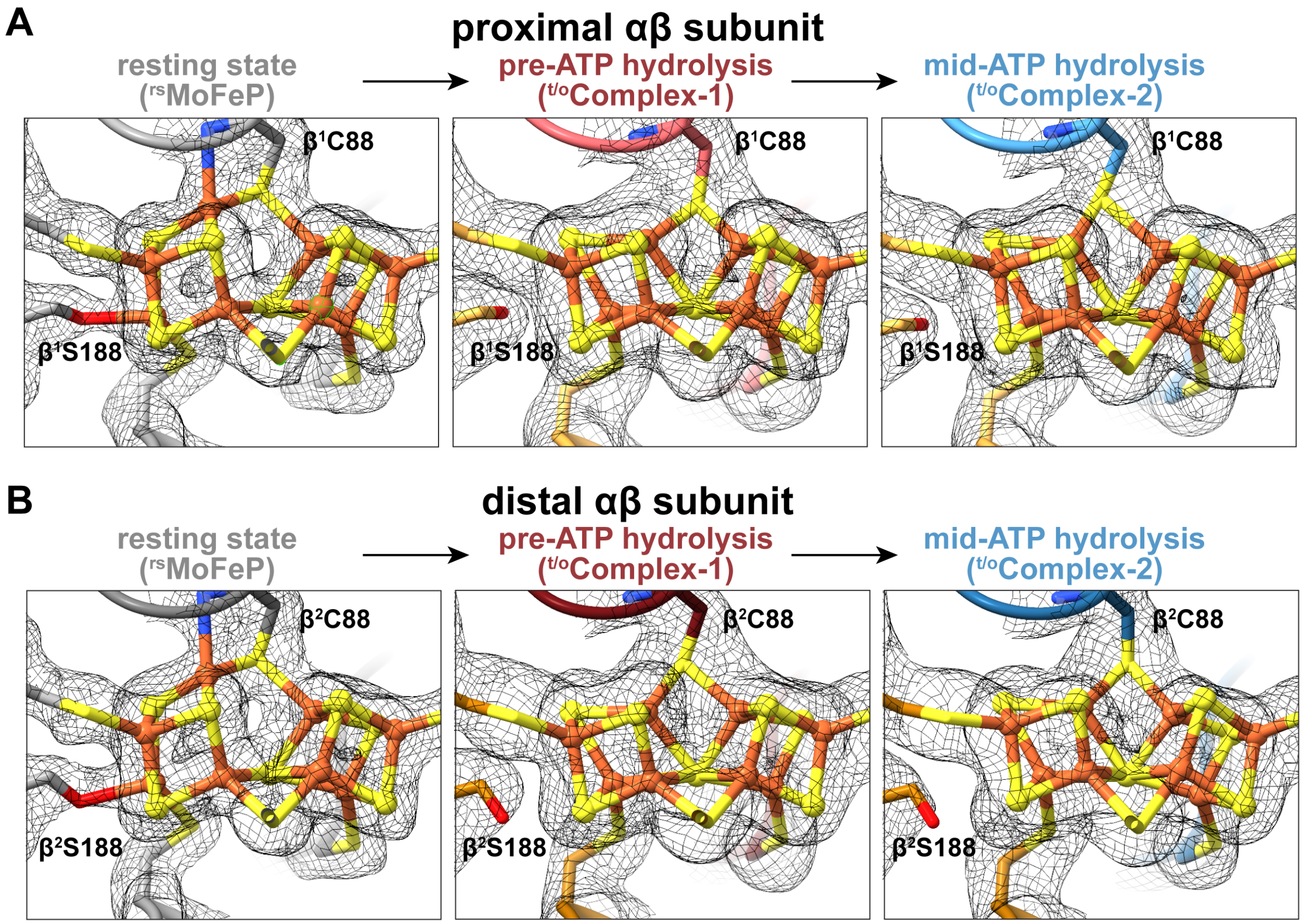


**Figure S8. P-clusters of rsMoFeP, $^{t/o}$Complex-1, and $^{t/o}$Complex-2. (A,B)** Views of the P-cluster and P-cluster ligands in the proximal (A) and distal (B) $\alpha\beta$ halves of MoFeP in rsMoFeP (gray), $^{t/o}$Complex-1 (maroon), and $^{t/o}$Complex-2 (blue) structures. CryoEM maps for each individual structure are contoured at the same level and shown as a gray mesh.

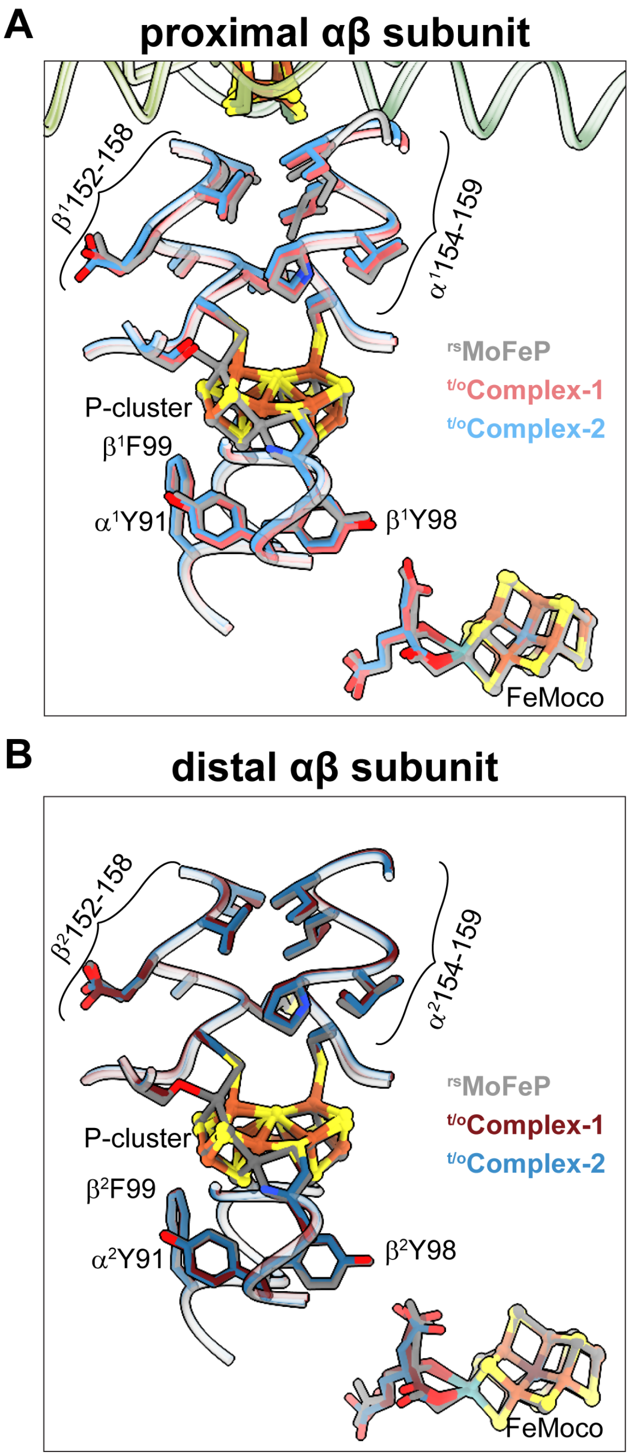


**Figure S9. Overlay of core residues along the ET pathway leading from the MoFeP surface to FeMoco. (A,B)** Overlay key residues residing between the [4Fe:4S] cluster and the P-cluster, and between the P-cluster and FeMoco in rsMoFeP (gray), $^{t/o}$Complex-1 (maroon), and $^{t/o}$Complex-2 (blue) structures. The proximal (A) and distal (B) $\alpha\beta$ halves of MoFeP are shown. $\gamma$100's helices of FeP are shown in green in (A).

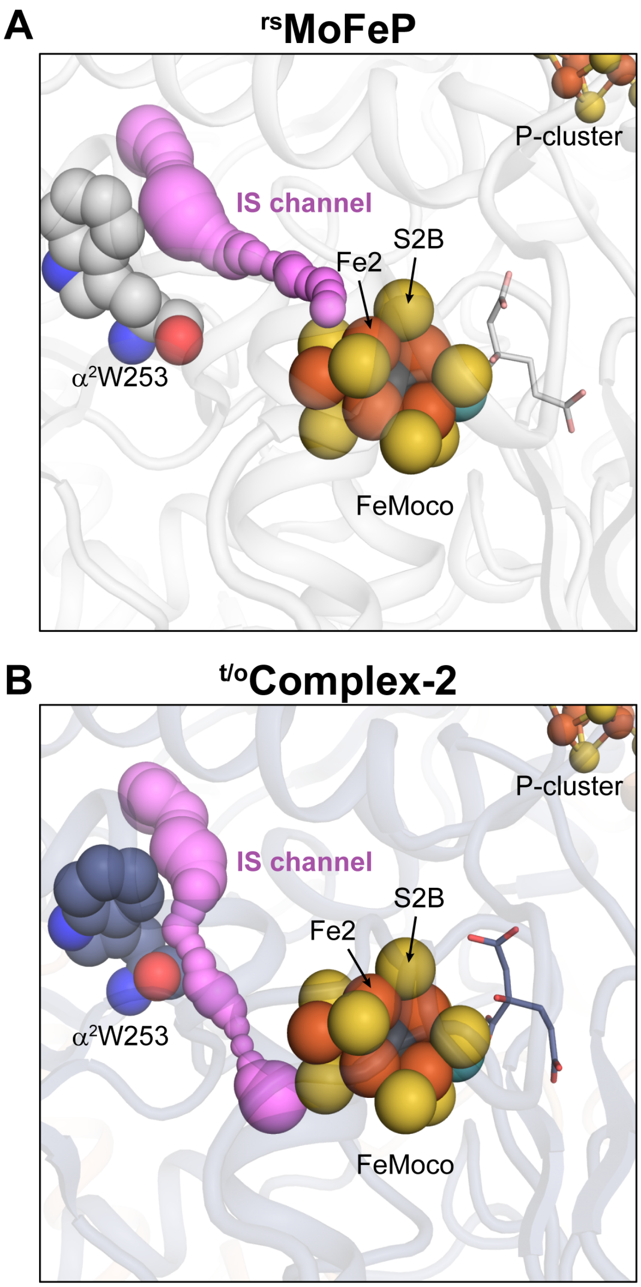


**Figure S10. Diversion of the IS channel during turnover. (A,B)** The proposed substrate pathway, termed the IS channel (12) is shown in pink and was calculated using the software CAVER (13) in PyMOL (14). (A) In rsMoFeP, the IS channel leads from the surface of MoFeP to the proposed catalytic face of FeMoco. (B) Conformational changes during turnover, as shown in the case of $^{t/o}$Complex-2, divert the IS channel to a different face FeMoco.

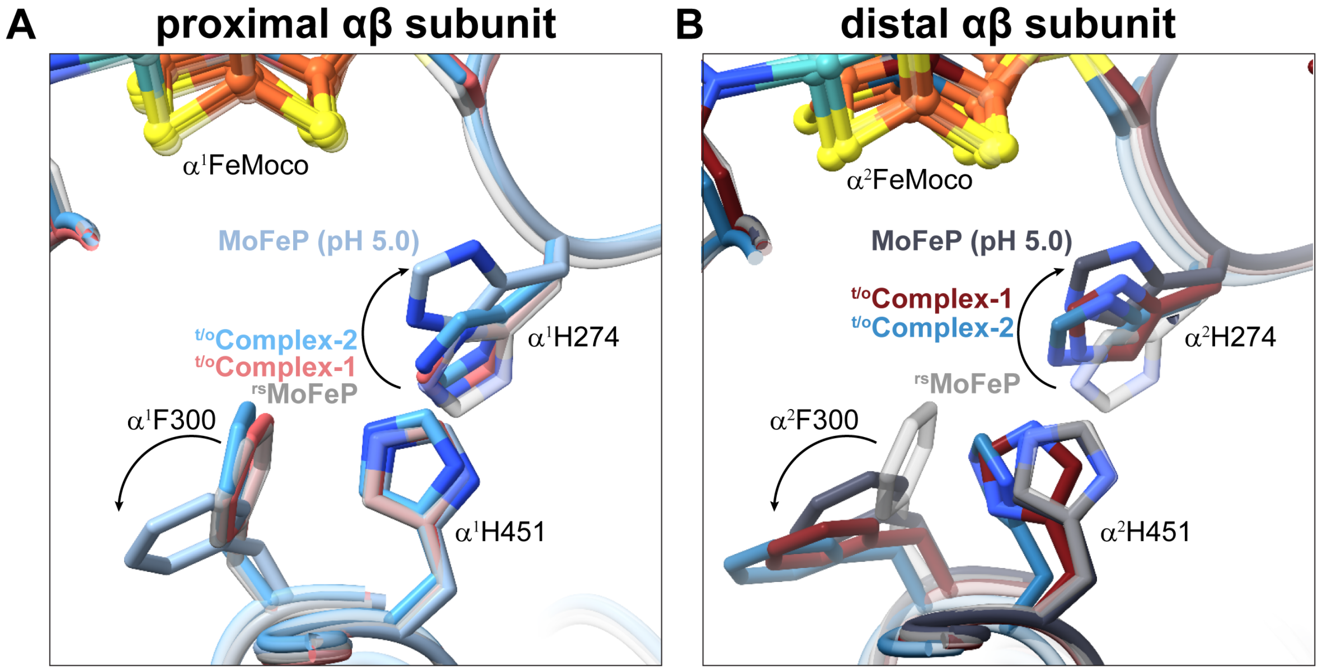


**Figure S11. Overlay of $\alpha$His274, $\alpha$Phe300 and $\alpha$His451 conformations observed in nitrogenase complexes under turnover and previously determined low-pH crystal structure of MoFeP. (A,B)** Overlay of the conformationally altered residues in rsMoFeP (transparent gray), $^{t/o}$Complex-1 (maroon), $^{t/o}$Complex-2 (blue), and the crystal structure of MoFeP determined at pH 5.0 (sky blue/slate PDB ID: 5VQ4) structures in both the proximal (A) and distal (B) $\alpha\beta$ halves of MoFeP.

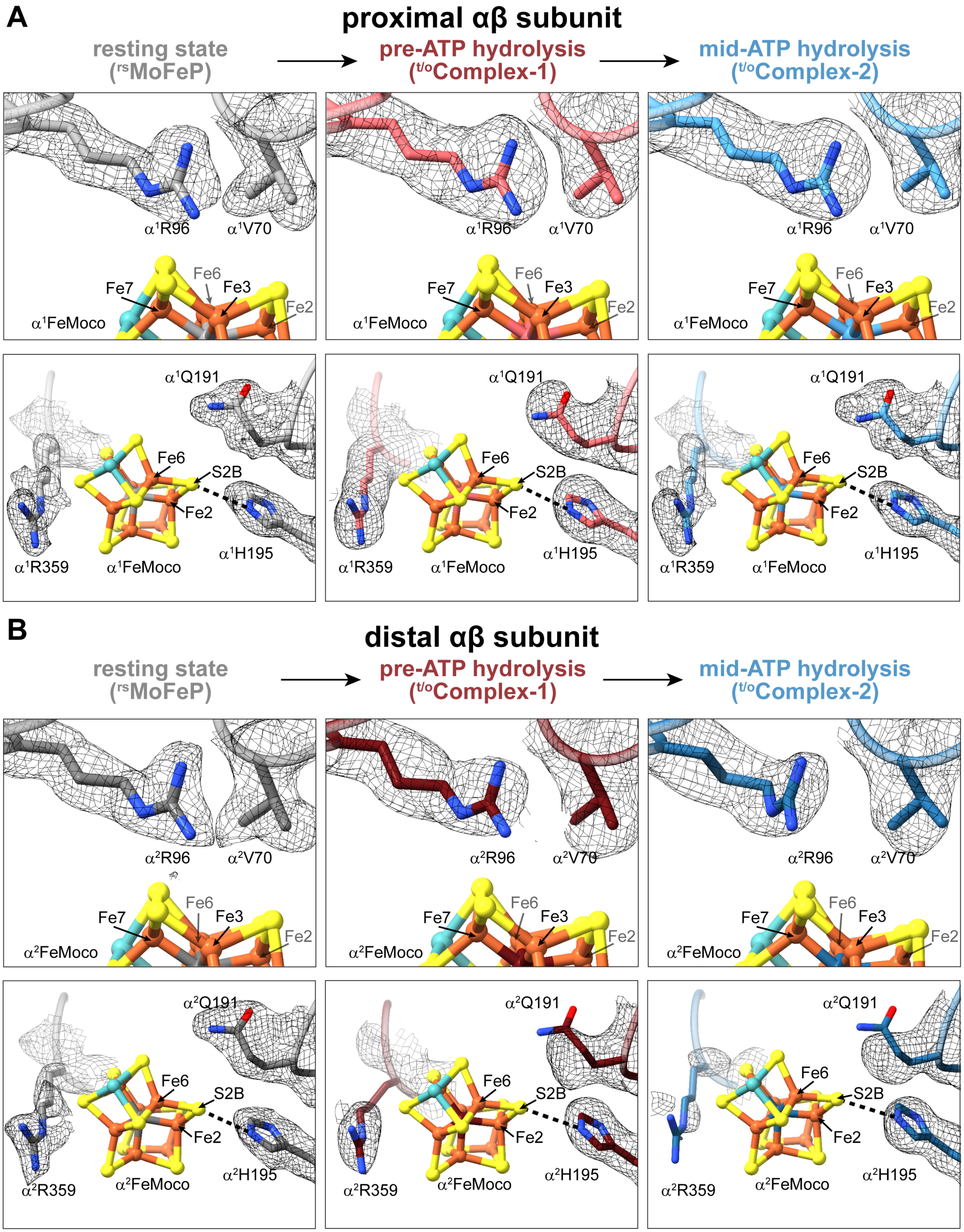


**Figure S12. The FeMoco environment in the cryoEM structures. (A, B)** Views of FeMoco and the nearby residues $\alpha$V70, $\alpha$R96, $\alpha$Q191, $\alpha$H195, and $\alpha$R359 in the proximal (A) and distal (B) $\alpha\beta$ halves of MoFeP in rsMoFeP (gray), $^{t/o}$Complex-1 (maroon), and $^{t/o}$Complex-2 (blue) structures. CryoEM maps for each individual structure are contoured at the same level.

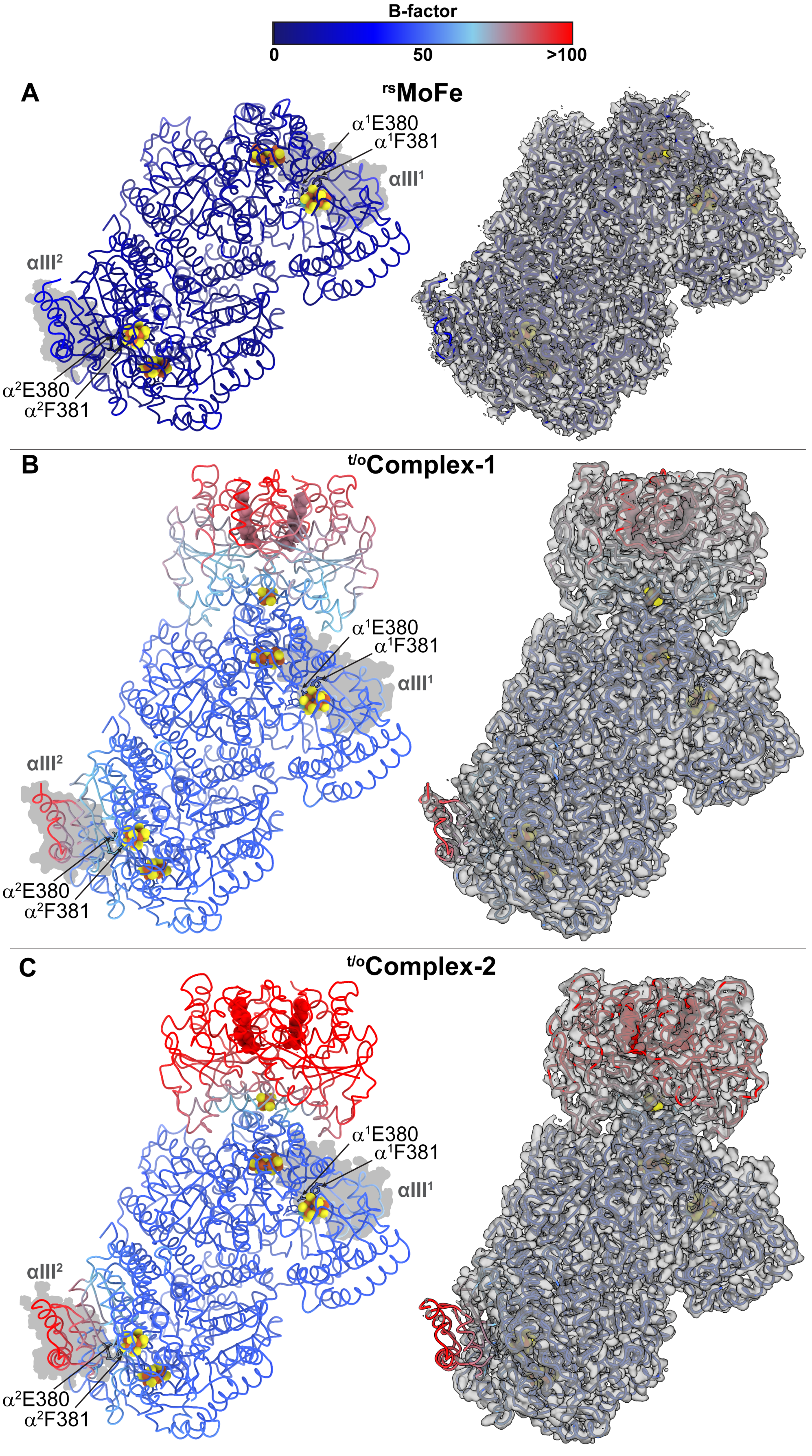


**Figure S13. t/oComplex-1, and t/oComplex-2 colored by B-factor and comparison of $\alpha$III densities.** **(A)** rsMoFeP, (B) t/oComplex-1, and (C) t/oComplex-2, colored by B-factor (left) and comparison of cryoEM map densities (right).

| | #1 $^{t/o}$Complex-1 (EMDB-26760) (PDB-7UT8) | #1 $^{t/o}$Complex-2 (EMDB-26763) (PDB-7UT9) |
|---|---|---|
| **Data Collection** | Titan Krios G3i K3 BioQuantum | |
| Magnification | 130kx | |
| Voltage (kV) | 300 | |
| Spherical Aberration (mm) | 2.7 | |
| Electron Exposure ($e^-/Å^2$) | 65 | |
| Defocus range (µm) | -1.1 to -2 | |
| Pixel size (Å, Physical/Digital) | 0.835 | |
| Energy Filter Slit Width (eV) | 20 | |
| Movies | 2580 | |
| **Map Statistics and Post-Processing** | $^{t/o}$Complex-1 | $^{t/o}$Complex-2 |
| Symmetry imposed | $C_1$ | $C_1$ |
| Map Resolution (Å) | 2.28 | 2.29 |
| Local resolution range for 75% of voxels | 2.731 | 2.577 |
| Local resolution range (model) | 1.897 - 21.602 | 1.843 - 22.574 |
| Map sharpening *B* factor ($Å^2$) | 47.5 | 47.3 |
| Map sharpening method | DeepEMhancer | DeepEMhancer |
| 3-D FSC values<br>X<br>Y<br>Z | <br>2.78<br>2.78<br>2.78 | <br>2.66<br>3.28<br>2.66 |
| **Model Statistics and Validation** | $^{t/o}$Complex-1 | $^{t/o}$Complex-2 |
| Model composition<br>Non-hydrogen atoms<br>Protein residues<br>Nucleic acids<br>Ligands | <br>20194<br>20026<br>0<br>168 | <br>20182<br>20018<br>0<br>164 |
| *B*-factors ($Å^2$)<br>Protein/Nucleic acid atoms<br>Ligands/non-protein atoms | <br>58.63<br>68.84 | <br>63.87<br>79.58 |
| R.M.S deviations<br>Bond lengths (Å)<br>Bond angles (˚) | <br>0.006<br>1.475 | <br>0.006<br>1.496 |
| MolProbity Score | 1.81 | 2.04 |
| MolProbity Clashscore | 7.80 | 9.36 |
| CaBLAM (% outliers) | 1.79 | 1.95 |
| Rotamer outliers (#,%) | 1.48 | 2.27 |
| Cis peptides (#, %) | 8.1, 0.3% | 8.1, 0.3% |
| Ramachandran Plot<br>Favored (%)<br>Allowed (%)<br>Disallowed (%) | <br>96.28<br>3.48<br>0.24 | <br>96.00<br>3.76<br>0.24 |
| EM Ringer score | 4.03 | 3.98 |
| Map/Model FSC (0.5) | 2.95 | 2.84 |

**Table S1.** CryoEM data collection and refinement statistics of *Azotobacter vinelandii* $^{t/o}$Complex-1 and $^{t/o}$Complex-2.

| | **#1 $^{rs}$MoFe $C_1$ (EMDB-26756) (PDB-7UT6)** | **#1 $^{rs}$MoFe $C_2$ (EMDB-26757) (PDB-7UT7)** |
|---|---|---|
| **Data Collection** | Titan Krios G3 K2 BioContinuum | |
| Magnification | 165kx | |
| Voltage (kV) | 300 | |
| Spherical Aberration (mm) | 2.7 | |
| Electron Exposure ($e^-/Å^2$) | 65 | |
| Defocus range (µm) | -0.5 to -1.5 | |
| Pixel size (Å, Physical/Digital) | 0.815 | |
| Energy Filter Slit Width (eV) | 10 | |
| Movies | 1460 (session 1)<br>2313 (session 2) | |
| **Map Statistics and Post-Processing** | | |
| Symmetry imposed | $C_1$ | $C_2$ |
| Map Resolution (Å) | 1.91 | 1.81 |
| Local resolution range for 75% of voxels | 5.299 | 2.883 |
| Local resolution range (model) | 1.826 - 30.347 | 1.826- 29.563 |
| Map sharpening *B* factor ($Å^2$) | 37.9 | 39.8 |
| Map sharpening method | DeepEMhancer | DeepEMhancer |
| 3-D FSC values<br>X<br>Y<br>Z | <br>2.47<br>2.78<br>2.37 | <br>1.93<br>1.94<br>1.84 |
| **Model Statistics and Validation** | | |
| Model composition<br>Non-hydrogen atoms<br>Protein residues<br>Nucleic acids<br>Ligands | <br>16142<br>15677<br>0<br>96<br>369 | <br>16142<br>15677<br>0<br>96<br>369 |
| R.M.S deviations<br>Bond lengths (Å)<br>Bond angles (˚) | <br>0.004<br>0.998 | <br>0.004<br>0.998 |
| MolProbity Score | 1.29 | 1.29 |
| MolProbity Clashscore | 4.56 | 4.56 |
| CaBLAM (% outliers) | 0.76 | 0.76 |
| Rotamer outliers (#,%) | 0.91 | 0.91 |
| Cis peptides (#, %) | 8, 0.4% | 8, 0.4% |
| Ramachandran Plot<br>Favored (%)<br>Allowed (%)<br>Disallowed (%) | <br>97.74<br>2.26<br>0.00 | <br>97.74<br>2.26<br>0.00 |
| EM Ringer score | 6.13 | 7.11 |
| Map/Model FSC (0.5) | 2.17 | 2.07 |

**Table S2.** CryoEM data collection and refinement statistics of *Azotobacter vinelandii* rsMoFeP.

| | **#1 $BeF_x$ trapped complex (EMDB-26764) (PDB-7UTA)** | |
|---|---|---|
| **Data Collection** | Titan Krios G3 K2 BioContinuum | |
| Magnification | 165kx | |
| Voltage (kV) | 300 | |
| Spherical Aberration (mm) | 2.7 | |
| Electron Exposure ($e^-/Å^2$) | 65 | |
| Defocus range (µm) | -0.5 to -1.5 | |
| Pixel size (Å, Physical/Digital) | 0.815 | |
| Energy Filter Slit Width (eV) | 10 | |
| Movies (Tilt (°)) | 1460 | 0 |
| | 2313 | 0 |
| | 2211 | -15 |
| | 2099 | -25 |
| **Map Statistics and Post-Processing** | 2:1 $BeF_x$ | |
| Symmetry imposed | $C_1$ | |
| Map Resolution (Å) | 2.40 | |
| Local resolution range for 75% of voxels | 2.883 | |
| Local resolution range (model) | 1.826- 29.563 | |
| Map sharpening *B* factor ($Å^2$) | 39.8 | |
| Map sharpening method | DeepEMhancer | |
| 3-D FSC values<br>X<br>Y<br>Z | <br>3.17<br>2.84<br>2.95 | |
| **Model Statistics and Validation** | | |
| Model composition<br>Non-hydrogen atoms<br>Protein residues<br>Nucleic acids<br>Ligands | <br>24304<br>24077<br>0<br>227 | |
| *B*-factors ($Å^2$)<br>Protein/Nucleic acid atoms<br>Ligands/non-protein atoms | <br>47.27<br>58.79 | |
| R.M.S deviations<br>Bond lengths (Å)<br>Bond angles (˚) | <br>0.011<br>1.137 | |
| MolProbity Score | 3.20 | |
| MolProbity Clashscore | 18.38 | |
| CaBLAM (% outliers) | 0.59 | |
| Rotamer outliers (#,%) | 13.95 | |
| Cis peptides (#, %) | 8, 0.3% | |
| Ramachandran Plot<br>Favored (%)<br>Allowed (%)<br>Disallowed (%) | <br>89.53<br>8.11<br>2.36 | |
| EM Ringer score | 3.67 | |
| Map/Model FSC (0.5) | 2.87 | |

**Table S3.** CryoEM data collection and refinement statistics of *Azotobacter vinelandii* $BeF_X$ trapped complex.

| structure | MoFeP (3u7q) | nucleotide-free complex (2afh) | AMPPCP-bound (4wzb) | ADP-bound (2afi) | AMPPCP/ADP-bound (4wza) | ADP.AlF$_x$ (1m34) | N-species-bound MoFeP (6ug0) | CO-bound MoFeP (4tkv) | $^{rs}$MoFeP | $^{t/o}$Complex-1 | $^{t/o}$Complex-2 |
|---|---|---|---|---|---|---|---|---|---|---|---|
| **nucleotide-free complex (2afh)** | 0.365 | | | | | | | | | | |
| **AMPPCP-bound (4wzb)** | 0.323 | 0.291 | | | | | | | | | |
| **ADP-bound (2afi)** | 0.386 | 0.329 | 0.374 | | | | | | | | |
| **AMPPCP/ADP-bound (4wza)** | 0.241 | 0.246 | 0.196 | 0.337 | | | | | | | |
| **ADP.AlFx (1m34)** | 0.396 | 0.319 | 0.318 | 0.401 | 0.318 | | | | | | |
| **N-species-bound MoFeP (6ug0)** | 0.232 | 0.324 | 0.283 | 0.328 | 0.283 | 0.378 | | | | | |
| **CO-bound MoFeP (4tkv)** | 0.106 | 0.329 | 0.224 | 0.361 | 0.224 | 0.385 | 0.224 | | | | |
| **$^{rs}$MoFeP** | 0.324 | 0.313 | 0.311 | 0.320 | 0.302 | 0.322 | 0.310 | 0.321 | | | |
| **$^{t/o}$Complex-1** | 0.434 | 0.397 | 0.359 | 0.412 | 0.386 | 0.348 | 0.410 | 0.444 | 0.233 | | |
| **$^{t/o}$Complex-2** | 0.458 | 0.416 | 0.385 | 0.420 | 0.411 | 0.375 | 0.436 | 0.465 | 0.246 | 0.170 | |
| **BeF$_x$-bound** | 0.437 | 0.399 | 0.382 | 0.429 | 0.404 | 0.366 | 0.433 | 0.444 | 0.261 | 0.246 | 0.252 |



**Table S4.** Root-mean-square-deviations (RMSDs; based on C$\alpha$ positions) between various MoFeP structures.

| Structure (resolution / PDB ID) | average B for the entire MoFeP component (Å$^2$) | average B for αIII (chain A) (Å$^2$) | average B for αIII (chain C) (Å$^2$) |
|---|---|---|---|
| MoFeP with oxidized P-cluster (2.03 Å / 2 MIN) | 24.13 | 34.70 | 34.14 |
| MoFeP at pH 5.0 (2.30 Å / 5VQ4) | 24.65 | 45.58 | 42.73 |
| 1.0-Å resolution MoFeP (1.0 Å / 3U7Q) | 10.58 | 11.37 | 13.13 |
| F99YMoFeP with oxidized P-cluster (1.4 Å / 6O7M) | 17.08 | 22.62 | 23.09 |
| Nucleotide-free FeP-MoFeP complex (2.1 Å / 2AFH) | 22.58 | 32.58 | 32.49 |
| MgADP-bound FeP-MoFeP complex - molecule 1 in the asymmetric unit (3.1 Å / 2AFI) | 32.42 | 45.70 | 45.67 |
| MgADP-bound FeP-MoFeP complex - molecule 2 in the asymmetric unit (3.1 Å / 2AFI) | 33.44 | 46.89 | 46.53 |
| MgAMPPCP/MgADP-bound FeP-MoFeP complex (1.9 Å / 4WZA) | 23.93 | 37.40 | 31.48 |
| MgAMPPCP-bound FeP-MoFeP complex (2.3 Å / 4WZB) | 27.35 | 45.01 | 37.91 |
| Crosslinked FeP-MoFeP complex (3.2 Å / 1M1Y) | 52.60 | 57.09 | 68.55 |
| MgADP.AlFx -stabilized MoFeP-FeP complex – molecule 1 in the asymmetric unit (2.30 Å / 1M34) | 33.45 | 44.61 | 48.94 |
| MgADP.AlFx -stabilized MoFeP-FeP complex – molecule 2 in the asymmetric unit (2.30 Å / 1M34) | 33.55 | 44.62 | 49.50 |

**Table S5.** Average B-factors for the MoFeP components and the $\alpha$III domains in various nitrogenase crystal structures.
